# Sexual Dimorphism in Age-Dependent Neurodegeneration After Mild Head Trauma in *Drosophila*: Unveiling the Adverse Impact of Female Reproductive Signaling

**DOI:** 10.1101/2024.03.06.583747

**Authors:** Changtian Ye, Ryan Ho, Kenneth H. Moberg, James Q. Zheng

**Affiliations:** Department of Cell Biology, Emory University School of Medicine, Atlanta, GA 30322, USA; College of Art and Science, Emory University, Atlanta, GA 30322, USA; Department of Neurology, Emory University School of Medicine, Atlanta, GA 30322, USA; Center for Neurodegenerative Diseases, Emory University School of Medicine, Atlanta, GA 30322, USA

**Keywords:** Brain Injury, Neurodegeneration, Aging, sex difference, fruit flies, mild TBI

## Abstract

Environmental insults, including mild head trauma, significantly increase the risk of neurodegeneration. However, it remains challenging to establish a causative connection between early-life exposure to mild head trauma and late-life emergence of neurodegenerative deficits, nor do we know how sex and age compound the outcome. Using a *Drosophila* model, we demonstrate that exposure to mild head trauma causes neurodegenerative conditions that emerge late in life and disproportionately affect females. Increasing age-at-injury further exacerbates this effect in a sexually dimorphic manner. We further identify Sex Peptide (SP) signaling as a key factor in female susceptibility to post-injury brain deficits. RNA sequencing highlights a reduction in innate immune defense transcripts specifically in mated females during late life. Our findings establish a causal relationship between early head trauma and late-life neurodegeneration, emphasizing sex differences in injury response and the impact of age-at-injury. Finally, our findings reveal that reproductive signaling adversely impacts female response to mild head insults and elevates vulnerability to late-life neurodegeneration.

## INTRODUCTION

Neurodegenerative disorders affect millions of people worldwide, and this number is expected to increase substantially with rising life expectancy and population growth. Major efforts have been directed towards understanding the etiology of neurodegenerative conditions, but the challenge remains to elucidate molecular mechanisms underlying the onset and progression of neurodegeneration. Recent studies have identified multiple genetic mutations and risk factors associated with dementia and neurodegeneration, but additional causes are likely involved ^1-4^. Aging is a major risk factor for many neurodegenerative disorders and plays a key role in disease emergence and progression ^5,6^. Other significant risk factors include environmental insults that can evoke neuropathological processes, which may be latent at first but surface with aging to disrupt various aspects of brain structure and function, leading to neurodegenerative conditions ^7,8^. Physical blows to the head are known to cause symptoms related to immediate brain injury, but even seemingly innocuous head impacts are strongly linked to brain dysfunction and degeneration later in life ^8-10^. It is thus crucial to understand the aberrant processes triggered by these mild physical insults that may be initially “hidden” but can give rise to later neurodegenerative conditions. Furthermore, sex differences have been documented in many degenerative diseases, including chronic brain degeneration following mild traumatic brain injury ^11-13^. How sex and age contribute to an individual’s response to mild brain disturbances and modify the risk for neurodegeneration remains to be fully understood.

The fruit fly, *Drosophila* melanogaster, represents an excellent and tractable model organism for dissecting fundamental disease mechanisms, including neurodegeneration ^14-17^. Importantly, the fruit fly’s relatively short lifespan enables longitudinal interrogation of disease progression from the initial trigger to the late-life emergence of neurodegenerative conditions such as Alzheimer’s disease (AD), amyotrophic lateral sclerosis (ALS), and frontotemporal dementia (FTD) ^17-21^. Recently, several *Drosophila* head injury models have been developed that recapitulate key findings from traumatic brain injury (TBI) in humans and other preclinical models, revealing significant insights into the potential molecular and genetic underpinnings of injury responses ^22-28^. We recently developed a novel *Drosophila* model (HIFLI: <u>H</u>eadfirst <u>I</u>mpact <u>FLy</u> <u>I</u>njury) in which mild repetitive head-specific impacts can be delivered to multiple awake and unrestricted adult flies of both sexes ^28^. In this study, we utilized the HIFLI model to deliver a milder version of repetitive head trauma, which eliminates potentially confounding effects from injury-induced death and enables us to interrogate sex and age-at-injury contributions to the emergence of brain deficits throughout the entire lifespan. Our data show that exposure to this milder form of head trauma elicits minimal acute deficits but causes profound brain-associated behavioral deficits and brain degeneration that only emerge later in life. These late-life neurodegenerative conditions are further exacerbated by increasing age-at-injury and disproportionately elevated in mated females. We further identify that Sex Peptide signaling involved in female reproduction plays a key role in elevated female neurodegeneration after mild head trauma. Finally, RNA sequencing data suggest that the chronic suppression of innate immune defense networks in mated females may mediate the elevated vulnerability to neurodegeneration after mild head trauma. Together, our results further establish *Drosophila* as an excellent model system to investigate life-long neurodegenerative conditions, highlight that sex and age significantly contribute to trauma recovery and outcome, and support the notion that physical insults to the head, even very mild ones, pose a major threat for brain health.

## METHODS

### Fly Husbandry

Flies were maintained at 25 °C, with 60% humidity on a 12-h:12-h light: dark cycle and kept in vials containing fresh fly media consisting of cornmeal, yeast, molasses, agar, and p-hydroxy-benzoic acid methyl ester. Vials were changed every 2-3 days to ensure fresh food and minimize flies getting stuck to wet food. The following stocks were used: *w^1118^* (BDSC, 5905), *neuronal synaptobrevin-GAL4* (*nSyb-GAL4)* (generously provided by Dr. T. Ngo from the Rubin Lab at Janelia Research Campus), SP null mutant line (0325/TM3, *Sb, ry*), SP deficiency line (Δ130/TM3, *Sb, ry*) ^29^ (both generously shared by Dr. M. Wolfner at Cornell University), *SPSN-splitGAL4* ^30^ (generously provided by Dr. K. Koh at Thomas Jefferson University), and *UAS-SPR RNAi* (BDSC, 66888). SP^0^ males (0325/ Δ130) were generated by crossing the SP null mutant line to the deficiency line. Pan-neuronal and SPSN-specific knockdown of SPR was achieved by crossing *UAS-SPR RNAi* females to *nSyb-GAL4* and *SPSN-splitGAL4* males, respectively. Unless otherwise noted, flies used were progeny from *nSyb-GAL4* males crossed to *w^1118^* females. Virgin females and males were collected within 8 hours of eclosion and kept in separate vials. For mated flies, 1 day after collection, female and male flies were given 24 hours to mate and separated again after. Flies were handled meticulously without CO_2_ after separating by sex to negate anesthetic effects.

### Injury paradigm and apparatus

Flies were injured in accordance with the protocol detailed in Behnke 2021 ^28^. Briefly, multiple flies were contained within a custom plastic injury vial designed to fit within an injury rig consisting of a cradle connected to a pulley system with a counterweight. To ensure consistency across injuries, each injury trial consisted of pulling the cradle downwards to the base of the injury rig, followed by lightly tamping the cradle three times for all the flies to fall to the bottom of the vial before releasing it for upward acceleration. When the cradle and injury vial stopped at the top, the momentum of the fruit flies continued to propel them upwards to the glass ceiling, where they sustained headfirst impacts. In this study, we modified the injury paradigm such that it consisted of 1 session of 15 impacts spaced 10 seconds apart (referred to as <u>v</u>ery <u>m</u>ild <u>H</u>ead <u>T</u>rauma, vmHT). Flies were injured at 3 days (D3_Inj_), 17 days (D17_Inj_), or 31 days post eclosion (D31_Inj_). Sham flies (Control) were placed into the injury vials for the duration of an injury session (2.5 minutes) but did not receive any injuries. For subsequent data analyses, we combined all sham groups within each experiment as we observed no behavioral or pathological differences between them.

A subset of flies was subject to an eye-penetrating injury as a positive control for TUNEL staining. Here, flies were anesthetized on a CO_2_ pad while a fly pin penetrated one side of the eye and into the brain. Flies were given one day to recover before fixation and subsequent immunofluorescent staining. Dead flies were excluded from subsequent analyses.

### Video-assisted startle-induced climbing assay

Startle-induced climbing behavior was assessed using a modified Negative Geotaxis Assay (mNGA). Climbing was assessed at the following timepoints after injury: 90 minutes, 1 day, 1 week, and weekly until D45. Flies were placed in plastic vials with a soft bottom consisting of 5% agar. Custom 3-D printed caps wrapped in parafilm were used to contain the flies. Up to four vials were assayed at the same time, using a custom 3-D Polylactic Acid (PLA) printed rig. A lightboard was placed behind the apparatus to enhance background contrast for the video recordings. For each trial, the rig was lightly tamped three times, and fly movement was recorded using a Panasonic HC-V800 digital video recorder at 60 frames per second. Three trials were conducted for each video, spaced 1 minute apart. Using FIJI, videos were cropped to fit individual vials, and then trimmed to the first 601 frames (10 sec) after flies fall to the bottom. The 10-second videos then underwent manual fly behavior tracking or idTracker.ai and subsequent analyses. All testing took place between ZT 3 and ZT 8 (ZT, Zeitgeber time; lights are turned on at ZT 0 and turned off at ZT 12) and testing occurred between 20 and 23 °C under laboratory lighting conditions.

Manual fly tracking was done using FIJI. Briefly, using the segmentation tool, the length of each vial was segmented into 10 equal bins and the number of flies in each bin was recorded at every second (every 60 frames), starting at the first frame. A Climbing Index for each second was calculated using the sum of the number of flies in each bin multiplied by the bin number and then divided by the total number of flies in the trial. Repeated measures ANOVA tests were used to compare the effects of the various ages-at-injury conditions on Climbing Indices for each second of the trials and post hoc tests were performed using the Bonferroni method.

Automated AI-powered tracking of fly negative geotaxis behavior was performed using idtracker.ai ^31^, a software utilizing a deep-learning algorithm that permits simultaneous positional tracking of multiple flies at high temporal resolution. 10-second NGA videos were processed in an Anaconda environment using a Dell workstation equipped with a NVIDIA Quadro RTX5000 graphic card. Tracking results were manually validated using the validate tool provided by idtracker.ai before generating fly coordinates in regard to the XY plane. Superimposed video clips with the speed tail for each fly were generated using the video tool from the idtracker.ai package. Python scripts based on Trajectorytools from idtracker.ai were used to determine the speed and directional orientation frame-by-frame in addition to climbing trajectories for each fly. Data was imported into Microsoft Excel and histogram plots were generated using Excel. Statistical analyses were performed using two-sample t-tests.

### Immunohistochemistry

Immunohistochemistry and image analyses were performed as described in our previous publication ^32^. Whole flies were fixed for 3 h in 4% paraformaldehyde (PFA) in phosphate-buffered saline (PBS) containing 0.5% Triton-X (PBS-T) at room temperature with nutation. Flies were next rinsed 4 times for 15 min each with PBS-T and brains were subsequently dissected in PBS. Brains were permeabilized overnight in 0.5% PBS-T at 4 °C with nutation then blocked in 5% normal goat serum (NGS) in PBS-Triton for 90 min at room temperature with nutation. Brains were stained with Hoechst 33342 solution (1:1000; Thermo Scientific™ 62249) and Alexa Fluor 594 (AF594) phalloidin (1:400, Invitrogen A12381). Regions devoid of Hoechst and phalloidin signal that are not associated with physiological structures such as the esophagus were considered vacuoles. For TUNEL staining, brains were incubated with the mixture of two solutions in the *In Situ* Cell Death Detection Kit, Fluorescein (Roche-11684795910) at 37 °C for one hour. For structural analyses of pigment dispersing factor (PDF) neurons and mushroom bodies, brains were incubated with 5% NGS and primary antibodies (1:100 DSHB mouse anti-PDF C7 or 1:50 mouse anti-Fasciclin II 1D4) in 0.5% PBS-T for 2 days at 4 °C with nutation. Following primary antibody staining, brains were then washed four times in 0.05% PBS-T and then incubated with 5% NGS and secondary antibody (1:400 Alexa 594 goat anti-mouse IgG, Invitrogen A11032) in 0.5% PBS-T for 2 days at 4 °C with nutation. Stained brains were rinsed 4 × 15 min with PBS-T, followed by one wash in PBS for 90 min and then mounted on glass slides within SecureSeal^TM^ Imaging Spacers (Grace Bio-Labs, Bend, Oregon, USA) containing SlowFade^TM^ Gold Antifade mounting medium (Life Technologies, Carlsbad, CA, USA S36937). Whole-brain imaging at 1 μm steps was performed using an Olympus FV1000MPE two-photon system equipped with a water immersion objective XLPLN25XWMP (working distance of 2 mm and N.A. of 1.05). For TUNEL staining, imaging was performed using a confocal microscope (Nikon C2+ confocal system based on a Ti2-E microscope) at 1 μm steps. Image analysis was performed using ImageJ (Fiji) software. The number and size of vacuoles were recorded and statistically assessed using non-parametric Wilcoxon rank sum tests and post hoc tests using the Bonferroni method.

### RNA extraction and sequencing

Injured and sham flies were injured at 3 days post-eclosion. At 1 day and 6 weeks post-injury, brains from males, wildtype-mated females, virgin females, and SP^0^-mated females were collected, resulting in 16 conditions, each condition containing three independent cohorts of n=40 flies. RNA from brain tissue was extracted by TRIzol^®^/Chloroform extraction (ThermoFisher, 15596026) and further clean-up with the Zymo DNA Clean & Concentrator-5 kit (D4004) with on column DNase treatment (ThermoFisher, AM1907). Once obtained, RNA samples were transferred on dry ice to Admera Health for library preparation and sequencing. Total RNA integrity was checked by BioAnalyzer. QIAseq FastSelect rRNA Fly + NEB Ultra II Directional kit was used to deplete rRNA and prepare stranded cDNA libraries. Samples were sequenced on NextSeq High Output Flow Cell (Illumina) for 150 cycles to generate a total of 60M paired-ended, 101 base-pair reads.

### Bioinformatic analyses

Following sequencing, raw FASTQ reads passing QC filter (Q > 30) were obtained from BaseSpace. Analyses were conducted on the Galaxy web platform using the public server at usegalaxy.org ^33^ and then on R on a local machine. Quality control on the data was performed using FastQC ^34^, and data was subsequently mapped using RNA STAR ^35^ and the Ensembl ^36^ gene annotation for *Drosophila melanogaster* reference genome (dm6). Mapping results were examined using IGV ^37^, IdxStats from the Samtools suite ^38^, and Gene Body Coverage tool and Read Distribution from the RSeQC tool suite ^39^. Mapped reads were then assigned to exons/genes and tallied using featureCounts ^40^. To enable inter-sample read count comparisons, count normalization and differential expression analysis was conducted using DESeq2 ^41^ with models that incorporate sex, age, and injury conditions. Functional enrichment analyses of any differentially expressed genes were conducted using enrichGO from clusterProfiler 4.0 ^42^. Manual searches and annotations of genes were performed using FlyBase ^43^.

### Data reporting and statistical analysis

No statistical methods were used to determine sample sizes but are consistent with sample sizes of those generally employed in the field. All experiments contained 3 biological replicates, unless otherwise noted. All flies in each vial were administered the same treatment regimen. For each experiment, the experimental and control flies were collected, treated, and tested at the same time. For behavioral and pathological data, statistical analyses for comparisons were as described above. All statistical analyses were performed using R packages (R v4.2.2, [rstatix]). p values are indicated as follows: ***p<0.001; **p<0.01; *p<0.05.

## RESULTS

### Exposure to a milder form of head impacts elicited no mortality but resulted in age-related acute sensorimotor deficits

The HIFLI system delivers head-first impacts to multiple adult flies at ∼5m/s ^28^. Our previous study employing the HIFLI model delivered two separate sessions of 15 iterative impacts, with a one-day interval between the two sessions. Young flies exposed to this injury paradigm exhibited a small percentage of acute mortality and a significant reduction in lifespan, but flies subjected to 1 session exhibited little to no acute mortality ^28^. To enable lifelong investigation of the age-at-injury as a risk factor for neurodegeneration with minimal deaths, we reduced the severity of our paradigm to only one session of 15 impacts, hereafter referred to as vmHT (very mild Head Trauma). We then quantified survival, sensorimotor deficits, and brain pathology at various timepoints after exposure to vmHT (Figure 1A). As reported in our previous publication, around 50% of flies exhibit concussive-like behaviors immediately after receiving headfirst impacts, including uncoordinated movements, freezing, and loss of postural stability, but all flies recovered fully within the first five minutes ^28^. Importantly, vmHT had no significant effects on both short- and long-term survival when delivered on post-eclosion Day 3, Day 17, or Day 31 (Figure 1B), confirming the very mild nature of our injury paradigm.

**Figure 1.**
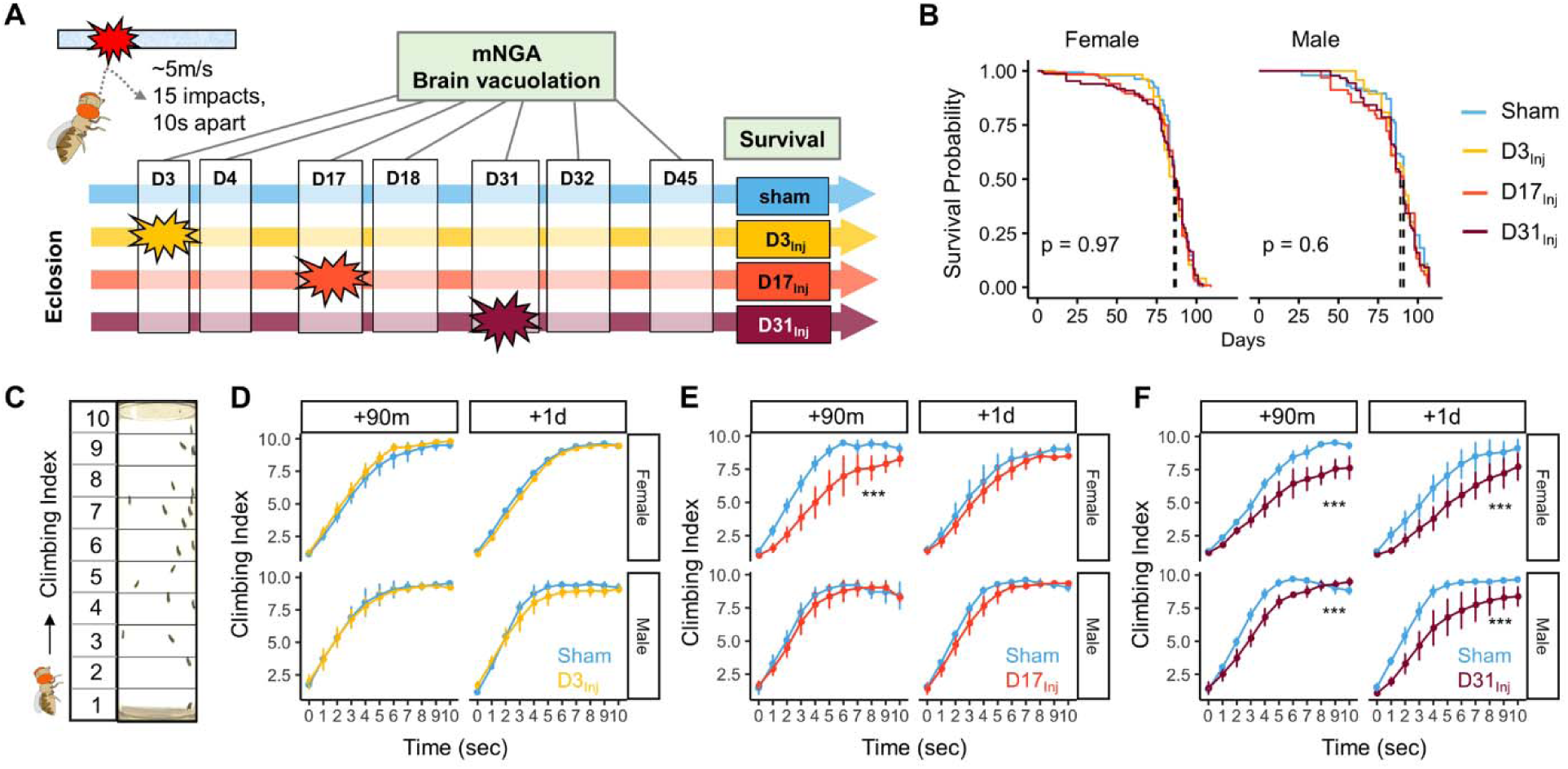
Exposure of adult fruit flies to a very mild form of repetitive head impacts at different ages elicits minimal acute effects. (**A**) Schematic illustration of the experimental design. Male and female flies were exposed to vmHT on D3, D17, or D31 post-eclosion (denoted as D3_Inj_, D17_Inj_, and D31_Inj_), whereas the sham groups did not receive vmHT. Sensorimotor behavior and brain pathology were assessed at 90 min and 1 day after vmHT exposure using mNGA and imaging-based quantification of vacuolization. (**B**) Survival curves showing no significant alteration in lifespan among different age-at-injury groups and between sexes. Kaplan-Meier p-values were determined using the Mantel-Cox log rank test with Bonferroni correction. Total N=871, n>50 flies in each condition. (**C**) Schematic diagram depicting mNGA where the Climbing Index (CI) at each second of the 10 second trial was calculated using the number of flies in each of the ten height bins (see Methods for details). (**D-F**) CI plots depicting acute effects of vmHT on the climbing behavior of D3_Inj_, D17_Inj_, and D31_Inj_ cohorts. **(D)** Both male and female D3_Inj_ flies showed no deficits in climbing at 90 min and 1-day post-injury. **(E)** Only female D17_Inj_ flies exhibited climbing deficits at 90 min post-injury (*** p=5.75e-08) but they recovered 1-day post-injury. **(F)** Both male and female D31_Inj_ flies exhibited marked decline in climbing ability at both time points (p values: 5.36e-08 and 6.2e-06 for female +90m and +1d; 0.00033 and 1.91e-07 for male +90m and +1d). A total of 9 videos were used for each condition: 3 experimental repeats of 3 trials, number of flies in each video n≥ 10. Error bar: ±se. Repeated measures ANOVAs were conducted to examine the effects of injury on climbing indices at each second.

Sensorimotor behaviors were assessed using a modified Negative Geotaxis Assay (mNGA, see Methods). We performed mNGA at 90 minutes and 1-day (1dpi) following vmHT exposure to examine acute injury effects and potential short-term recovery. Using fly positional data, we calculated a Climbing Index (CI) for each second of the 10 second trial (Figure 1C). Male and female flies injured on day 3 post eclosion (D3_Inj_) did not exhibit any acute deficits in climbing behavior (Figure 1D), though this was not the case for flies injured on day 17 (D17_Inj_) or day 31 (D31_Inj_). We noted a significant reduction in CI in D17_Inj_ females at 90 minutes post-injury, but not in males of the same age-cohort (Figure 1E). These females appeared to have mostly recovered by 24 hours post-injury (Figure 1E). In comparison, both male and female D31_Inj_ cohorts exhibited a substantial reduction in CI at 90 minutes as well as 1 dpi (Figure 1F), indicating age-related acute sensorimotor deficits after vmHT exposure and impaired recovery. These results are consistent with the notion that aging worsens post-head injury recovery ^44,45^ and that older populations are at a higher risk of developing negative consequences following mild head injury ^46-48^.

We next examined the short-term effects of vmHT on brain structure using a previously described protocol to stain whole fly brains and to detect vacuoles ^32^. Vacuolation, as seen in *Drosophila* models of neurodegenerative diseases, is a marker for frank tissue degeneration in the fly brain ^49,50^. These roughly round lesions can range from 1 to 50 μm in diameter and are largely located within the neuropil. Though aging alone can increase the number and size of vacuoles in the brain, we demonstrate that vmHT did not generate any changes in vacuolation at 1 dpi in all injury groups (Figure 2 A-C). Examination of the mushroom body of the central nervous system and the pigment dispensing factor (PDF) neurons in the optical lobe of D3_Inj_ flies also found no obvious structural alterations shortly after vmHT exposure (Supplementary Figure S1). Therefore, the observed acute climbing deficits were unlikely caused by acute structural disruptions of the brain after vmHT. Additionally, we performed TUNEL staining at 1 dpi to investigate whether vmHT causes apoptosis within the brain. As a positive control, we subjected a group of flies to an eye-stabbing procedure aimed at producing a very severe injury capable of inducing apoptotic cell death within the brain ^51^. We found that brains from the eye-stabbing injury group clearly displayed TUNEL-positive signals near primary injury regions, validating our TUNEL stain (Supplementary Figure S2). However, TUNEL staining was faint in the brains of both the shams and vmHT-exposed groups. Consistently, we did not observe any exoskeletal or eye damage immediately following injuries, nor did we observe any retinal degeneration and pseudopupil loss at the chronic stage of these flies. These data further support the mild nature of our head trauma model and suggest that acute sensorimotor deficits were not a direct result of neuronal death nor caused by gross brain disruptions.

**Figure 2.**
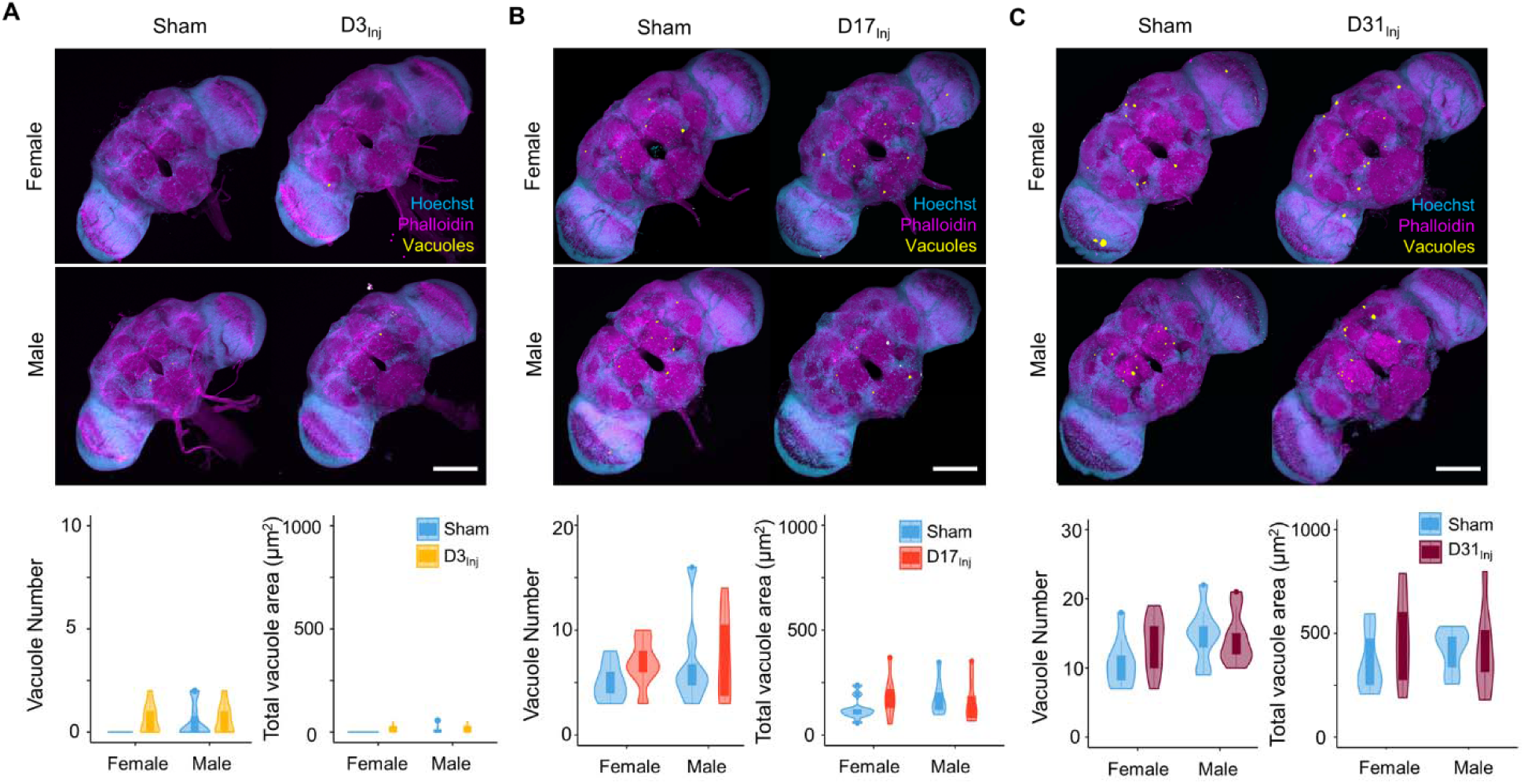
Exposure to vmHT does not acutely increase vacuole formation in the brain. (**A-C**) Two-photon whole brain imaging and quantification of vacuoles in D3_Inj_ (A), D17_Inj_ (B), and D31_Inj_ (C) flies. Top panels: representative z-projected whole-brain images of different injury groups with their respective sham brains of both sexes. Vacuoles are highlighted by yellow color. Scale bar = 100 μm. Bottom: violin plots and boxplots represent quantification of vacuole number and total vacuole area in each condition. Boxplots whiskers correspond to the maximum 1.5 interquartile range. Two experimental replicates resulting in n>10 brains for all conditions. Stats: non-parametric Wilcoxon rank sum tests, p>0.05.

#### Exposure to vmHT caused delayed onset of neurodegenerative conditions

Physical insults to the head, even very mild cases, represent a risk factor for the development of neurodegenerative conditions later in life. While exposure to vmHT did not generate obvious short-term brain-associated deficits in our younger cohorts (D3_Inj_ and D17_Inj_), we hypothesized that they could develop neurodegenerative conditions late in life. We quantified sensorimotor behavior and brain vacuolation for sham, D3_Inj_, D17_Inj_, and D31_Inj_ cohorts on day 45 post-eclosion (D45). In this age-matched comparison, we found that vmHT exposure decreased CI regardless of age-at-injury, though injury at older ages induced greater sensorimotor decline (Figure 3A). Importantly, sensorimotor deficits were sexually dimorphic as female flies appeared to perform worse than males in the same injury condition. To better visualize the sex differences, we plotted the accumulated CI (summation of CI values over the 10 second trial) and normalized the value to that of the respective sham groups (Figure 3B). In the D3_Inj_ cohort, only female flies exhibited significant climbing deficits when compared to the sham, whereas D17_Inj_ cohort females had a stronger decline in climbing ability than the males. Here, male flies seemed to be more resilient to vmHT when exposed at younger ages. When injured on D31, however, both male and female flies exhibited similarly profound sensorimotor deficits when assessed on D45.

**Figure 3.**
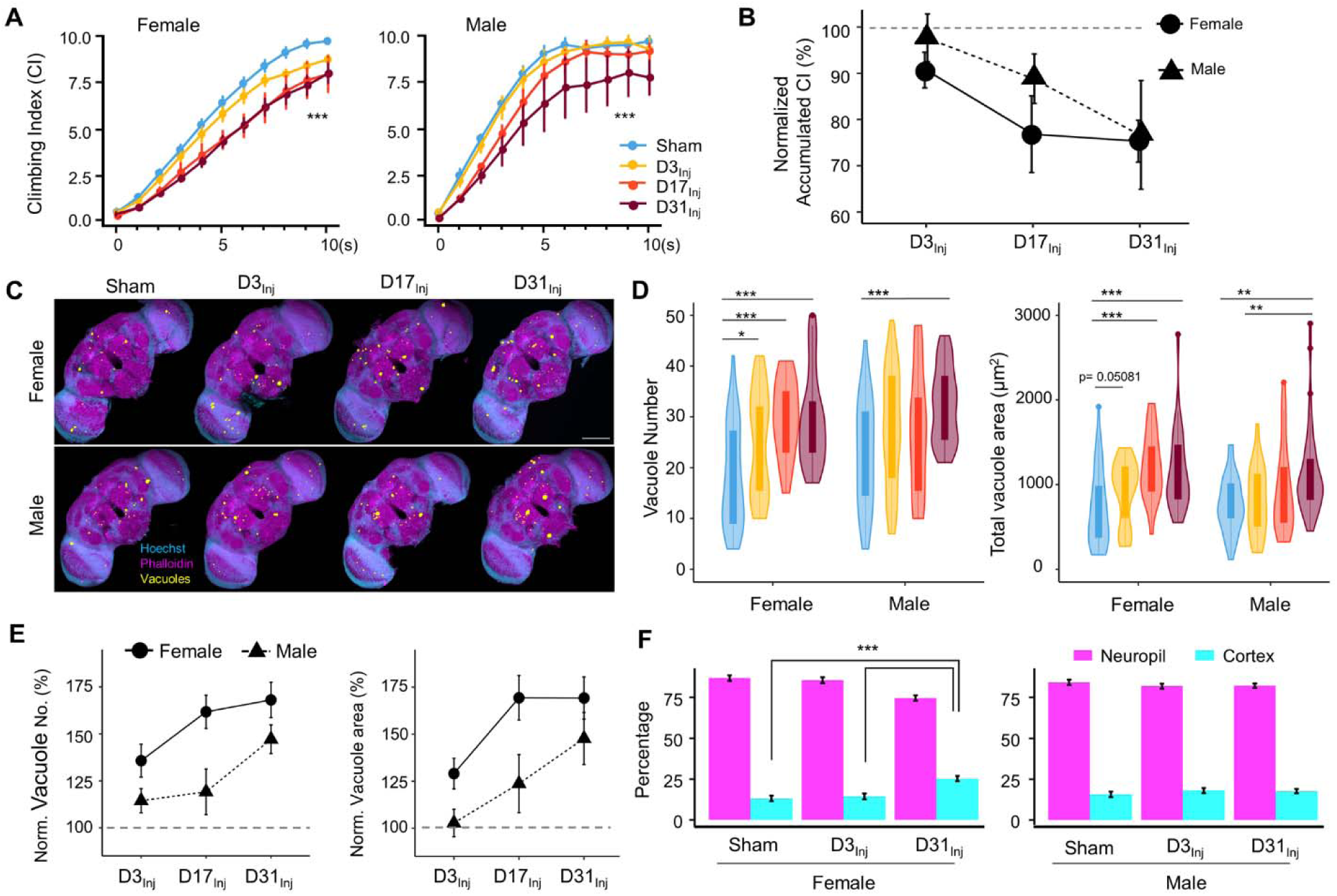
Exposure to vmHT results in late-life brain deficits and neurodegeneration. **(A)** Exposure to vmHT at various ages altered negative geotaxis behavior when assessed on D45. A total of 9 videos were used for each condition: 3 experimental repeats of 3 trials, number of flies in each trial n≥ 10. Error bar: ±se. Repeated measures ANOVAs and Bonferroni post hoc tests were conducted to examine the effects of injury on climbing indices at each second. Overall, vmHT exposure reduced CI for females (*** p=2e-16) and males (*** p=1.2e-11). See Supplementary Data S1 for p values from pairwise comparisons by injury conditions and time. **(B)** Sex differences in vmHT-induced climbing impairment are associated with age-at-injury. D3_Inj_ and D17_Inj_ females exhibit a stronger decline in normalized accumulated CI compared to males. D31_Inj_ females and males, on the other hand, suffered a similar substantial reduction in accumulated CI. Accumulated CI data of injury groups are normalized to their respective sham levels. **(C)** Representative z-projected whole images depicting vacuole formation in each condition (sex/ injury group). Scale bar = 100 μm. **(D)** Quantification of vacuole number and total vacuole area in each condition (Total N=352, n>30 in each condition). Here all sham groups were combined by sex. Boxplots whiskers correspond to the maximum 1.5 interquartile range. Statistics: non-parametric Wilcoxon rank sum tests. See Supplementary Data S1 for p values from pairwise comparisons by injury conditions. **(E)** Sex differences in the increase of brain vacuolation were partially dependent on age-at-injury. Vacuole number and total vacuole area of each injury condition were normalized to their respective sham groups. Overall, females exhibited a higher percentage of increase in vacuolation than males. **(F)** Quantification of the percentage of vacuoles in neuropil and cortex region of the brain in sham, D3_Inj_, and D31_Inj_ groups of both sexes. Data was from two independent trials of n=15 brains in each condition. Two sample t-tests were used to compare the percentage of vacuoles in neuropil and cortical region between each injury condition. D31_Inj_ females had significantly less percentage of vacuoles in the neuropil region compared to the sham group (*** p=2.19493e-6).

For female flies, significant increases in vacuole number and total area were observed for all three age-at-injury cohorts on D45 (Figure 3 C-D). For male flies, only the oldest injury cohort (D31_Inj_) showed a significant increase in vacuolation. When vacuole number and area were normalized to the respective sham groups and compared between sexes, we found that female brains exhibited a higher percentage of injury-induced increase in vacuolation overall, but particularly in younger injury cohorts D3_Inj_ and D17_Inj_ (Figure 3E).

In congruence with our previous finding that aging increases vacuolation, we found that flies aged to D45 exhibited a higher extent of brain vacuolation when compared to younger flies, even without injury. To further probe the origin of brain vacuolation, we examined the distribution of vacuoles by quantifying the percentage of vacuoles within the cell-body rich cortex and the axon-rich neuropil for sham, D3_Inj_, and D31_Inj_ flies on D45. We confirmed that regardless of injury condition, most vacuoles can be found in the neuropil and not in the cortex (Figure 3F). Interestingly, D31_Inj_ females have a higher percentage of cortical vacuoles when compared to sham and other injury cohorts, but males have a consistent neuropil to cortex ratio in all cohorts. Overall, this finding further suggests that neuronal death is not the major contributor of vacuole formation. Indeed, TUNEL staining at this time point (D45) did not yield significant signals for sham, D3_Inj_, D17_Inj_, or D31_Inj_ flies (Supplementary Figure S2D).

The above age-matched comparison highlighted that though vmHT exposure early in life has minimal acute effects, it can result in neurodegenerative conditions in advanced ages. While it inherently makes sense to compare flies at the same age (D45), different injury groups technically experienced different post-injury recovery times in such a comparison. Therefore, we also compared the groups by matching their recovery time of two weeks. We found that within the same timeframe, flies injured at younger ages exhibit a smaller decrease in sensorimotor abilities (Supplementary Figure S3A and B). This was true for both male and female flies, although sex difference existed in this 2-week comparison as well. Out of all injury groups, D3_Inj_ males were the only cohort of flies that displayed no change in climbing ability two weeks after vmHT exposure. On the other hand, all injured females, as well as D31_Inj_ males, exhibited an increase in vacuole formation (Supplementary Figure S3C-F). Overall, this recovery time-matched data suggests that aging accelerates the development of neurodegenerative conditions.

#### AI-assisted tracking revealed disruptions in fly speed and directional movement

The implementation of the Climbing Index in mNGA allows us to assess group climbing behavior and injury-induced deficits with an increased resolution in height (10 height bins) and time (every second). However, mNGA lacks the ability to reveal detailed information regarding sensorimotor behavior of individual flies, such as instantaneous speed and directional angle. Here, we took advantage of an AI-based tracking system, idtracker.ai ^31^, to track and analyze individual fly’s speed and direction at a temporal resolution of 1/60 sec (Figure 4A, see Supplementary video SV1). We were able to track 15-20 individual flies accurately and simultaneously through the entire course of an NGA trial (10 seconds). As shown in a representative tracked video from D45, female flies exposed to vmHT clearly showed an age-at-injury-dependent increase in the population of slow-moving flies, especially in the D31_Inj_ cohort (Supplementary Video SV1). In viewing NGA videos, it was also clear that though each trial is 10 seconds long, females took around 7 seconds to reach the top of the vial, whereas nearly all male flies reached the top of the vial by 4 seconds. Therefore, we elected to perform detailed quantitative analyses on climbing speeds and angles within the first 3 seconds after trial initiation, which better depicts the climbing responses to a startle stimulus.

**Figure 4.**
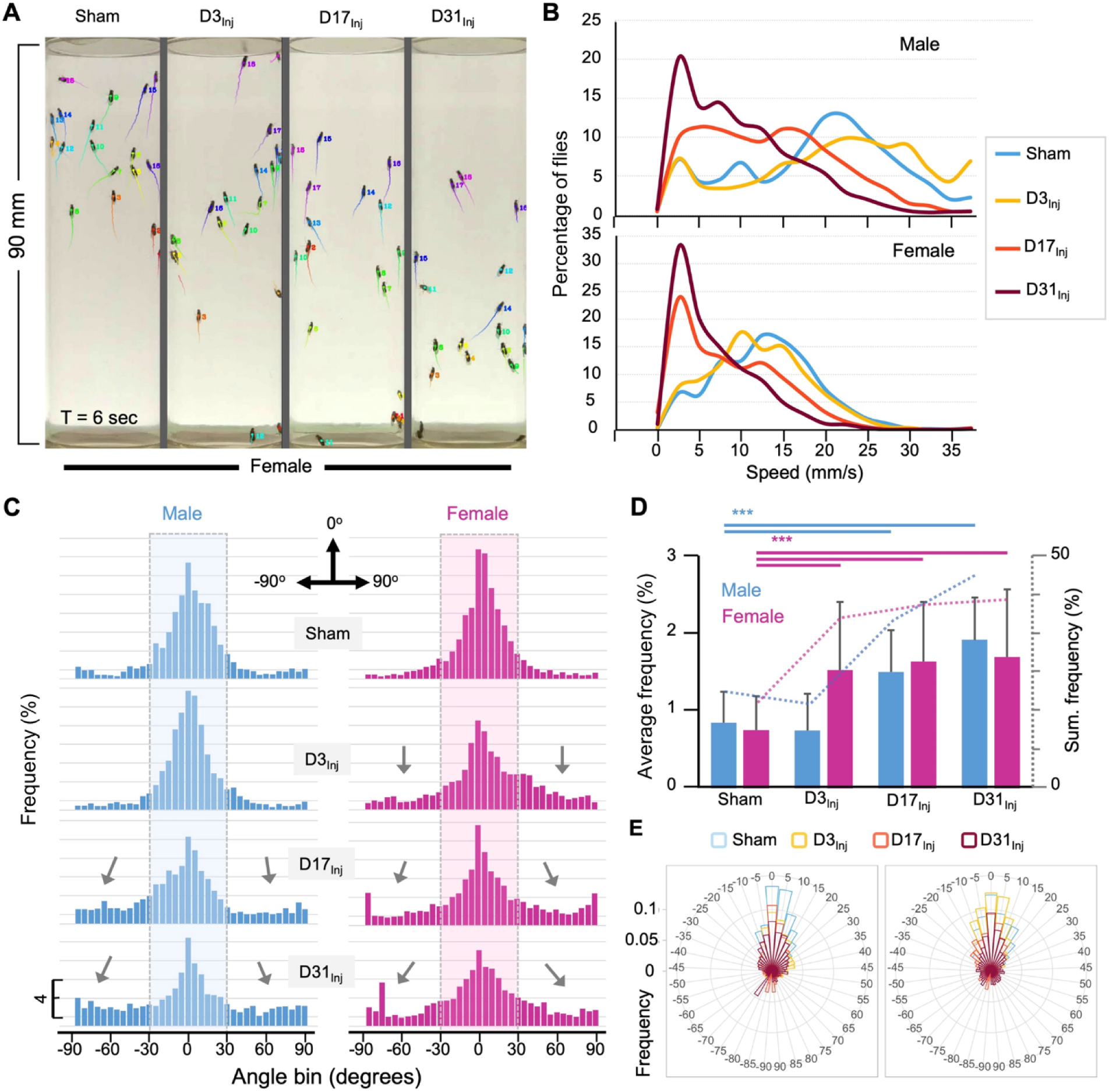
AI-assisted tracking and quantification of individual fly’s behavior provides insight into defects in the speed and direction of movement. (**A**) Representative snapshot of idtracker.ai-generated video at T=6 seconds. Videos used for analyses were from D45 flies (see Supplemental video SV1). Each trail represents fly movement speed calculated from the last 30 frames. Color is auto assigned to individual flies in each trial. (**B**) Histogram plots of the climbing speed of individual flies at a temporal resolution of 1/60 sec. The y-axis depicts the percentage of flies with a specific speed (x-axis). (**C**) Angular histograms showing the angle distribution of individual flies during the first 3 secs of mNGA trial. Highlighted sections represent normal fly directional orientation (between -30° and 30° with 0° as vertical). Arrows indicate the increased incidents of climbing angles outside the normal range. (**D**) Average percentages and accumulated percentages of flies with abnormal directional movement (<-30° or >30°) during the first three seconds of NGA trial. Two sample t tests, p<0.001, ***. (**E**) Rose plot depicting overall frequency distribution of fly orientation during the first three seconds of NGA trial.

Negative geotaxis represents a sensorimotor response that requires flies to first detect their position in respect to gravity and then move against the pull of gravity. Thus, injury-induced deficits could manifest in both the speed and angle of climbing. Speed analyses revealed that in both male and female flies, vmHT exposure resulted in a shift in the distribution of speed to slower speeds. Female flies exhibited a progressively larger shift from D3_Inj_, D17_Inj_, to D31_Inj_, and males only see this decrease in D17_Inj_ and D31_Inj_ cohorts, with D31_Inj_ flies exhibiting the slowest speed (Figure 4B). Next, we quantified the directionality of each fly at every frame of each trial and presented the data in angular histograms (Figure 4C). Here, vertical direction to the top is defined by an angle of 0°, whereas angles of -90° and 90° indicate that the fly is moving horizontally. We arbitrarily defined that flies moving with an angle between -30° and 30° are climbing normally in the vertical direction. vmHT exposure caused a significant portion of fly movement to fall outside the normal angular range, especially in flies injured at older ages (Figure 4C and D). Female flies in all injury groups exhibited significantly different and aberrant climbing angles when compared to the sham, whereas for males, only D17_Inj_ and D31_Inj_ groups exhibited significant changes to climbing angles (Figure 4E). Together, the impairments in movement speed and movement direction suggest that both sensory and motor pathways were disrupted by vmHT exposure.

#### Mating status determined female vulnerability to neurodegeneration following vmHT

Female flies are larger and about 30-40% heavier than males of the same age (Figure 5A). It is thus plausible that the elevated brain deficits observed in females were resulted from a larger impact force from vmHT. To test this possibility, we first reduced the impact force for female flies to match the impact force that the male flies experienced, which was accomplished by reducing the acceleration distance of the injury apparatus. We found that exposure to the reduced injury elicited similar behavioral deficits and brain pathology as exposure to regular vmHT (Supplementary Figure S4). Since virgin females are similar in size and mass to mated females (Figure 5A), we next investigated the long-term responses of virgin females to vmHT. First, we found that like mated females, injury did not affect survival in virgin females regardless of age-at-injury (Figure 5B). Next, we quantified sensorimotor behavior and vacuolation at D45, which is when we consistently see neurodegenerative conditions in mated females of different injury groups. Unlike the mated females, virgin females displayed little injury-induced climbing deficits nor showed significant changes in brain vacuolation (Figure 5C-E). Together, these data suggest that the difference in body size/mass between male and female flies is unlikely the cause for the observed vulnerability to late-life neurodegeneration in mated females. It appears that mating somehow insidiously alters the female biology, rendering it more vulnerable to developing neurodegeneration after injury.

**Figure 5.**
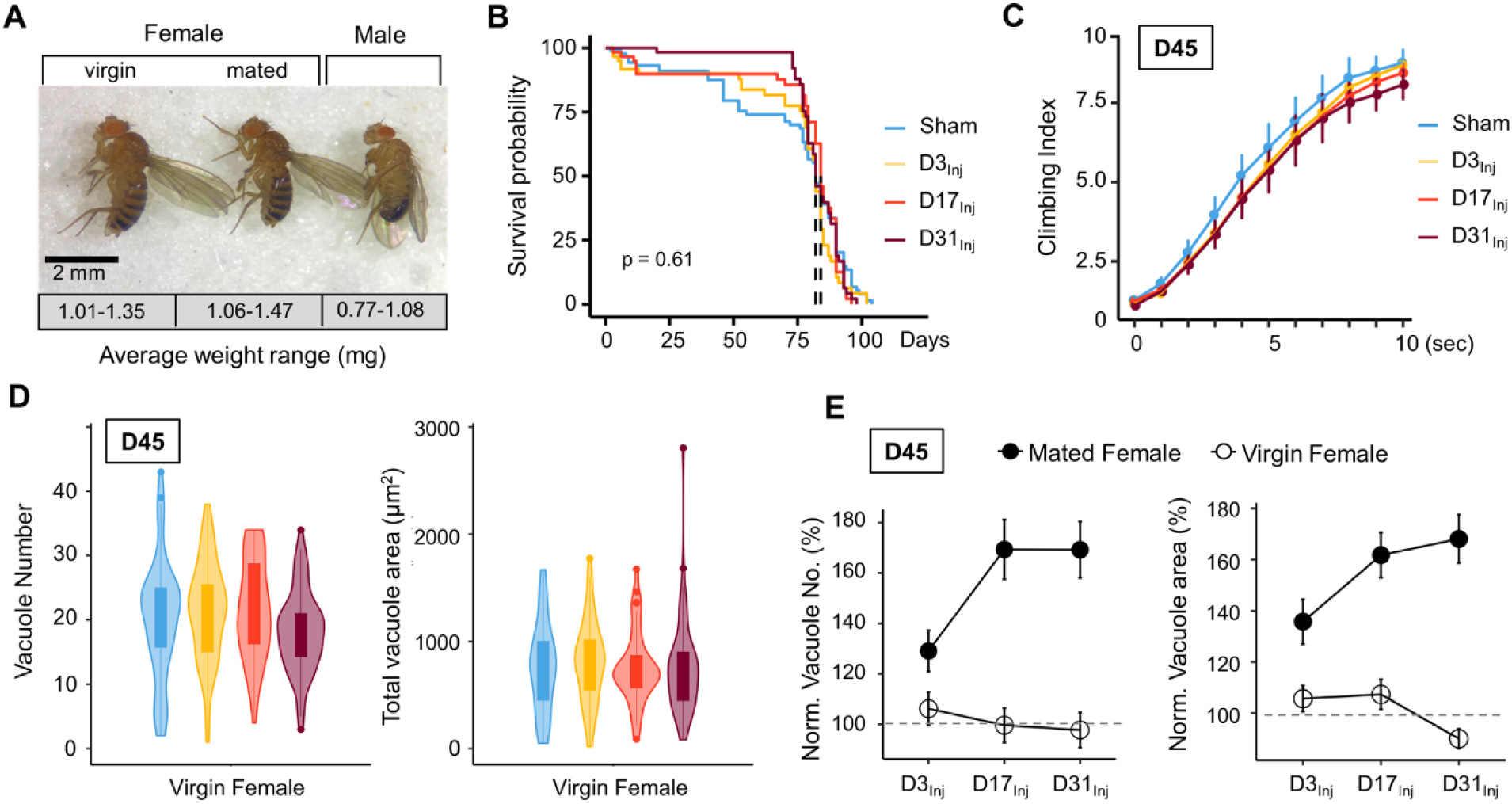
Virgin females do not exhibit neurodegenerative conditions regardless of age-at-injury. **(A)** Comparison of size and mass of virgin female, mated female, and male flies. 3 separate trials of n>100 in each condition. Total mass in each trial was averaged over the number of flies to generate a weight range for each condition. **(B)** vmHT resulted in no significant change to the lifespan of virgin females. Kaplan–Meier p-values were determined using the Mantel-Cox log rank test with Bonferroni correction. Total N=271, n>30 flies in each condition. **(C)** vmHT did not affect climbing behaviors of virgin females on D45 regardless of the age-at-injury. A total of 9 videos were used for each condition: 3 experimental repeats of 3 trials, number of flies in each trial n≥ 10. Repeated measures ANOVAs were conducted to examine the effects of injury on climbing indices at each second (p>0.05). **(D)** vmHT did not increase vacuole number or total vacuole area on D45 regardless of the age-at-injury. Total N= 224, n>25 for each condition. Statistics: non-parametric Wilcoxon rank sum tests. (**E**) Plots depicting differences in vacuole formation between virgin and mated females. Vacuole number and total vacuole area of each injury condition were normalized to the respective sham controls. Mated females exhibited much higher percentage of increase in vacuolation than virgin females.

#### Sex Peptide is responsible for female vulnerability

Mating is known to induce a wide array of behavioral and physiological changes in female flies, collectively known as post-mating responses. Female post-mating responses are primarily triggered by the male accessory protein Sex Peptide (SP) ^29,52^ binding to SP receptors (SPRs) ^53^ on a pair of Sex Peptide sensory neurons (SPSNs) ^54-56^ within the female reproductive tract. We next investigated the role of SP signaling in injury-induced neurodegenerative conditions in mated females. First, female flies were mated to SP^0^ males and then exposed to vmHT on D3, D13, or D31. mNGA and brain imaging on D45 revealed that these females behaved very similarly to the virgin females, with no lifespan reduction, sensorimotor deficits, or brain pathology due to vmHT at any age (Figure 6A). Next, pan-neuronal RNAi knockdown of SPRs in female flies yielded a similar result (Figure 6B). Like virgin females, these females exhibited no change in lifespan, negative geotaxis, or brain degeneration (Figure 6B). Finally, SPSN-specific RNAi knockdown of SPRs also abolished injury-associated deficits, regardless of age-at-injury (Figure 6C). It should be noted that the SPR RNAi line has been previously validated ^53^ and its effectiveness in SPR knockdown is evident in female flies as they exhibit dramatically reduced egg laying and, importantly, lack typical post-mating behaviors (such as the rejection of male flies after the initial mating) observed in the wild type-mated female flies. In fact, female flies with RNAi-mediated SPR knockdown behave identically to females mated with SP-null male flies, confirming the effective disruption of the SP-SPR signaling pathway. Together, these data suggest that SP signaling through the reproductive pathway is responsible for the observed vulnerability to injury-induced neurodegeneration in mated females.

**Figure 6.**
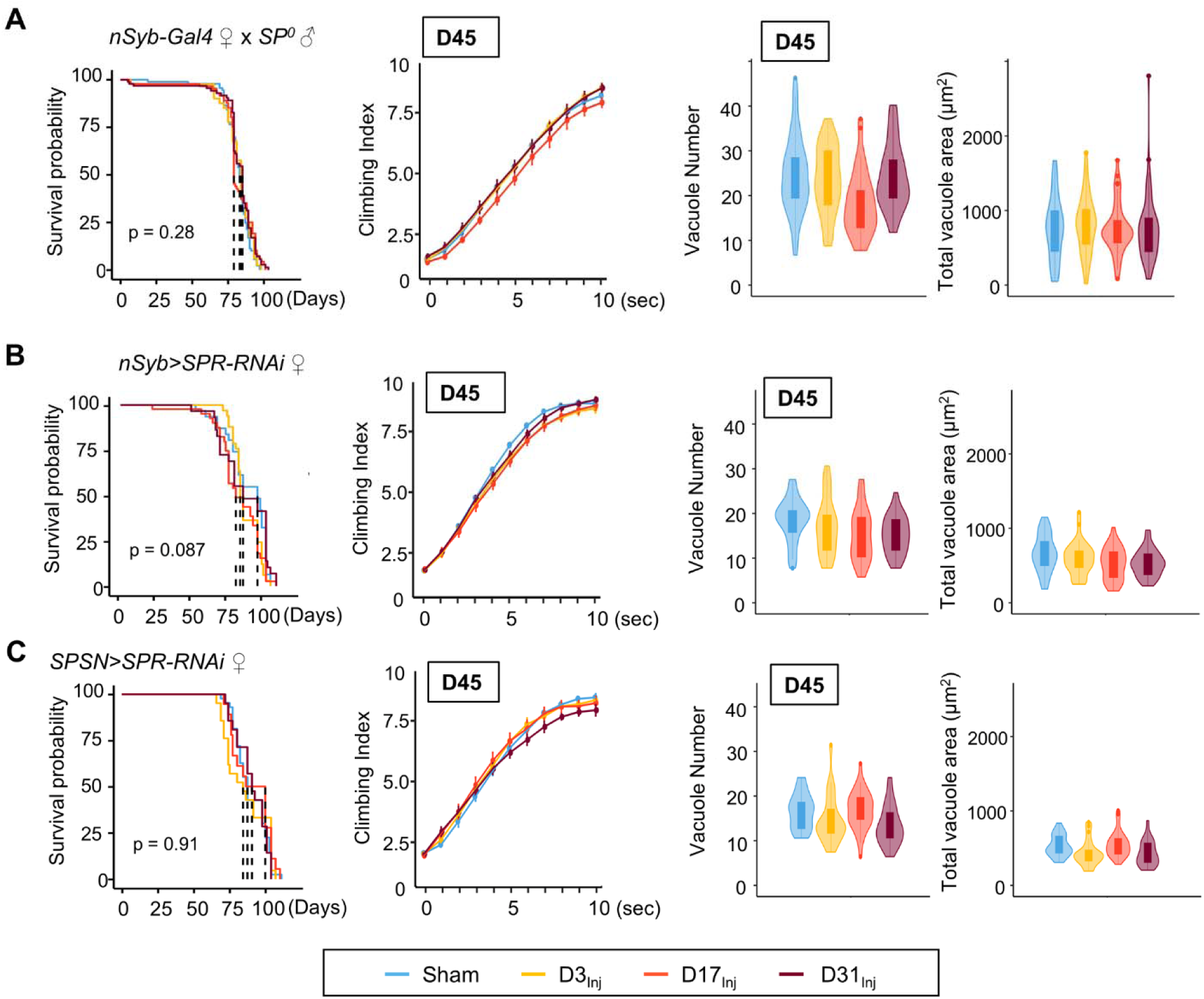
Eliminating SP signaling mitigates the emergence of neurodegenerative conditions in the female. (A-C) Survival curves, mNGA quantification, and vacuole quantification of female flies mated to SP^0^-males, females with pan-neuronal RNAi knockdown of SPR mated to wildtype males, and females with SPSN-specific knockdown of SPR mated to wildtype males (n>30 in each condition). No difference was detected between sham and injured groups in all three genotypes.

To further examine the relationship between SP signaling and injury-induced neurodegeneration, we performed a set of experiments in which virgin females were subjected to vmHT on D3, followed by mating with either wildtype or SP^0^ males for 24 hours on D10 (Figure 7A). We found that this post-injury mating scheme did not affect the lifespan of females that either mated with wildtype or SP^0^ males. Strikingly, post-injury mating with wild type males effectively reintroduced injury-induced phenotypes, including increased vacuole formation and decreased climbing. However, post-injury mating with SP^0^ males did not significantly alter the virgin injury response (Figure 7C-D). Together, these results suggest that in females, pathways activated by SP signaling and those elicited by vmHT likely interact and synergize to accelerate the progression of neurodegeneration.

**Figure 7.**
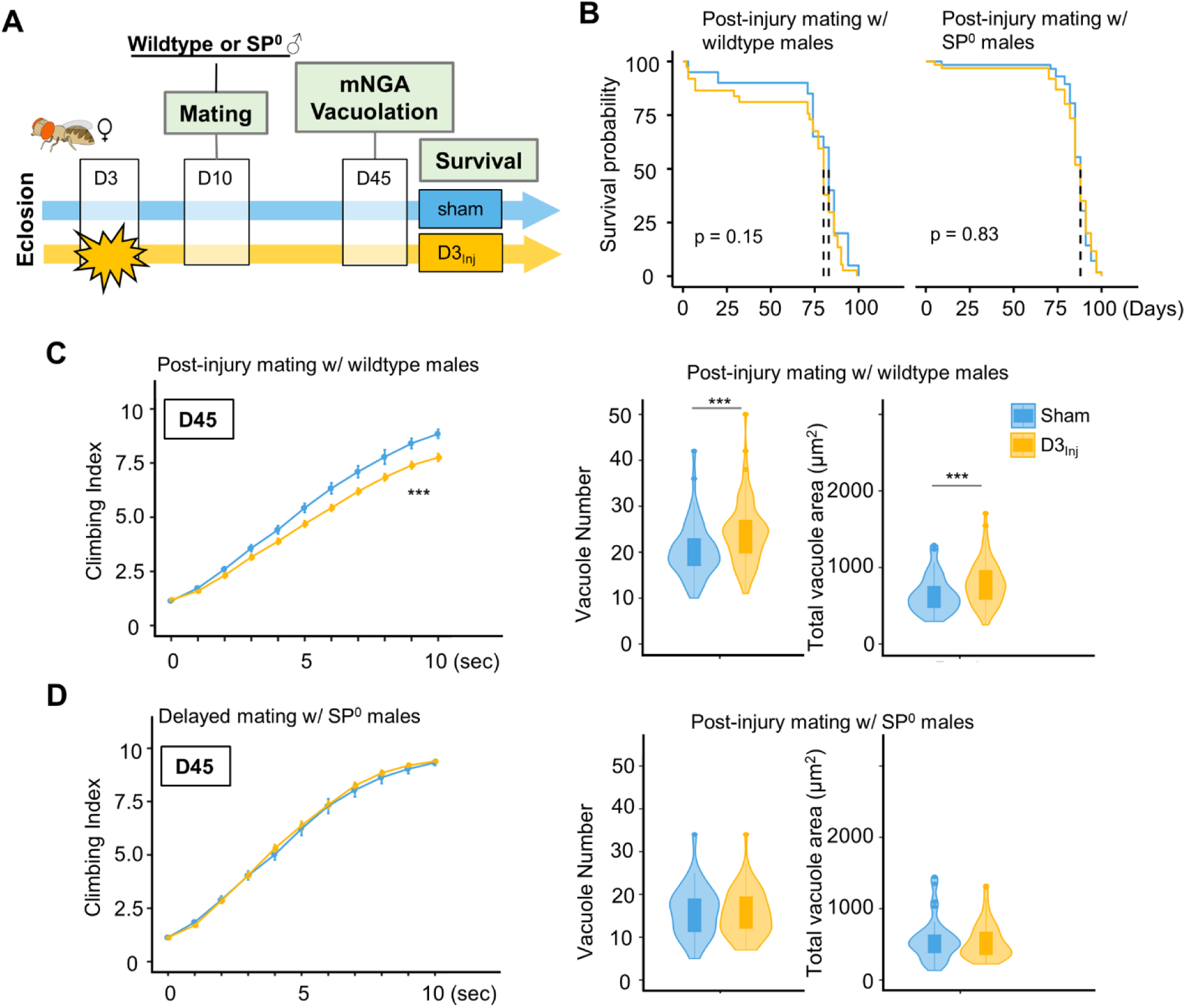
Introducing SP-signaling to pre-injured virgin females reinstates neurodegenerative phenotypes. **(A)** Diagram of mating schematic and relevant behavioral and pathological assays. Virgin female flies were subjected to vmHT on D3, whereas sham flies never receive an injury. On D10, these females were exposed to either wildtype or SP^0^ males for 24 hours to allow mating. mNGA and brain vacuolation were assessed on D45. **(B)** No change in the lifespan was observed for females that subjected to vmHT as virgin followed by post-injury mating with either male group. N= 182, n> 40 for each condition. **(C)** When assessed on D45, females subjected to post-injury mating with wildtype males exhibited significantly sensorimotor deficits (*** p=3.4e-15) and vacuole formation (vacuole number: p=0.0004536, vacuole area: p=0.0009563). A total of 9 NGA videos were used for each condition: 3 experimental repeats of 3 trials, number of flies in each trial n≥ 10. Error bar: ±se. Repeated measures ANOVAs and Bonferroni post hoc tests were conducted to examine the effects of injury on climbing indices at each second. For vacuole analyses, N= 149, n>30 in each condition. **(D)** When assessed on D45, females subjected to post-injury mating with SP^0^ male flies did not elicit sensorimotor deficits and vacuole formation. A total of 9 NGA videos were used for each condition: 3 experimental repeats of 3 trials, number of flies in each trial n≥ 10. For vacuole analyses, N= 105, n>20 in each condition.

#### RNA-seq analysis reveals sexually dimorphic responses to vmHT

To gain molecular insights into the sexually dimorphic responses to vmHT and the progressive development of late-life neurodegeneration, we conducted RNA sequencing on freshly dissected brain tissue from flies with and without vmHT exposure. Here, we focused on flies injured on day 3 after eclosion (D3_Inj_) since under this injury condition, only females mated with wildtype males exhibited behavioral and pathological deficits late in life (on D45, see Figure 3). Four distinct D3_Inj_ groups were chosen for RNA extraction and sequencing: wildtype males, females mated with wildtype males, virgin females, and females mated with SP^0^ males (Figure 8A). We assessed transcriptomic changes in the brain at two timepoints; 1-day post-injury (1dpi) for acute changes and 6-weeks post-injury (6wkpi) for chronic changes. Principal Component Analyses revealed that sex accounted for a large portion of variance in sequencing dataset, but the effects of vmHT were not immediately clear in either timepoints (Figure 8B). This observation reiterated the very mild nature of our injury paradigm. To further identify transcriptomic drivers of neurodegeneration after injury, we grouped the dataset by time post-injury, sex, and reproductive condition.

**Figure 8.**
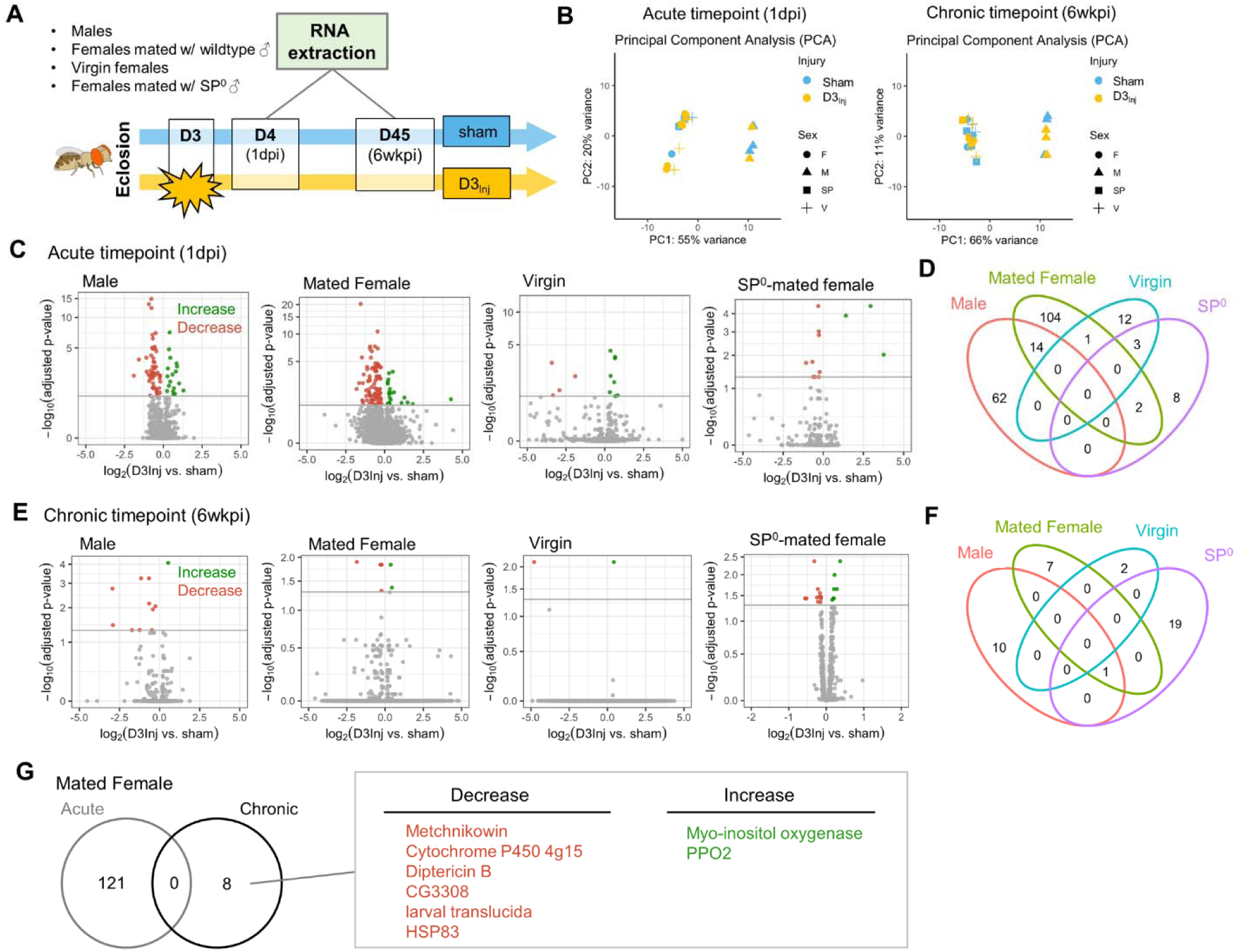
RNA sequencing reveals sexually dimorphic and reproductive status-dependent transcriptomic responses in D3_Inj_ fly brains. **(A)** Timeline and design of RNA-seq experiment. Sham and D3_Inj_ brains were collected at 1dpi and 6wkpi, in four different conditions (wildtype males, females mated with wildtype males, virgin females, and females mated with SP^0^ males), and in three biological replicates (n=40 brains per condition per replicate). Total sample number n= 48. **(B)** PCA analyses of samples at chronic and acute timepoints. **(C, E)** Volcano plots depicting gene expression at the acute and chronic timepoints. Upregulated genes are highlighted in green, downregulated genes are highlighted in red. **(D, F)** Venn diagram displaying the limited overlap of genes between the four conditions at the acute and chronic timepoints. **(G)** Venn diagram depicting no overlap between the acute and chronic gene changes in females mated to wildtype males. Upregulated genes are highlighted in green, downregulated genes are highlighted in red.

Compared to the respective sham groups, vmHT exposure resulted in significantly different sets of acute transcript changes between males and various female groups. D3_Inj_ induced 18 upregulated and 58 downregulated transcripts in male brains at 1dpi and 121 significant alterations (22 upregulated and 99 downregulated) in brains of wildtype-mated females (Figure 8C). However, we found a very few shared changes (14 genes) that are largely related to small molecule metabolic process (GO: 0044281) between male and female brains, suggesting sexually dimorphic responses to vmHT (Figure 8D, see Supplementary Data S2 for the complete list of differentially expressed genes). Interestingly, we observed a much smaller number of transcriptomic changes in virgin females and SP^0^-mated females (16 and 13, respectively) which minimally overlapped with injury-induced changes in male and wildtype-mated female brains. Overall, data from the acute timepoint suggest that the immediate response to vmHT is sexually dimorphic and dependent on reproductive status.

Next, we analyzed differential transcriptomic changes of sham and D3_Inj_ brains at the chronic timepoint (6wkpi). Strikingly, only a small number of transcripts were found to undergo significant changes in each condition (Figure 8E). In males, D3_Inj_ induced 1 upregulated and 10 downregulated gene transcripts on D45 when compared to the sham group. In wildtype-mated females, D3_Inj_ elicited 8 significant alterations (2 upregulated and 6 downregulated). Virgin female brains exhibited significant changes in only 2 transcripts (one upregulated and one downregulated), whereas SP^0^-mated female brains showed changes in 20 transcripts (15 upregulated and 5 downregulated) (for list of all differentially expressed genes, see Supplementary Data S3). We noted the lack of common injury-induced changes between all four conditions, except for one transcript shared between males, wildtype-mated females, and SP^0^-mated females (Figure 8F). This suggests that chronic transcriptomic profiles in response to vmHT were also sexually dimorphic and reproductive status dependent. Importantly, vmHT altered expression of 7 unique genes in wildtype-mated female brains. Three of these transcripts lie within immune system processes (GO:0002376), but the functions of the other four transcripts were not as clear. Specifically, mated females exhibited a decrease in the expression of two AMP genes, *Metchnikowin* and *Diptericin B*, both involved in the humoral immune response ^57^. We also observed an increase in the expression of *Prophenoloxidase 2*, whose activation is involved in the defense against pathogens ^58^. Furthermore, none of these genes were identified in the acute timepoint, suggesting that the altered expression of these genes may be involved in the late emergence of neurodegenerative conditions in mated females.

## DISCUSSION

An emerging body of evidence suggests that early exposure to repetitive mild head trauma increases the risk of developing neurodegenerative conditions later in life. However, the underlying mechanisms remain unclear. In this study, we take advantage of the short lifespan and powerful genetic tools of *Drosophila melanogaster*, in combination with our headfirst impact model ^28^, to investigate how early exposure to a very mild form of repetitive head trauma leads to late-life neurodegeneration. Previously, we showed that mild repetitive injuries can elicit degenerative phenotypes in both male and female flies, but female flies appear more vulnerable to the injuries ^28^. Our current study has further substantiated the previous findings by examining different age-at-injury cohorts and demonstrating their age-dependent and sexually dimorphic responses to a very mild form of repetitive head impacts. Furthermore, we discovered a strong link between SP signaling and injury-induced neurodegeneration in females. Compared to males, virgin females, and SP^0^-mated females, female flies mated to wildtype males were much more vulnerable to developing sensorimotor deficits and brain degeneration following injury exposure. This demonstrates, for the first time, the convergence and interaction of two supposedly separate pathways, namely the pathophysiological mechanisms elicited by mild head injury and the post-mating responses mediated by SP exposure. While activation of either one of these pathways was insufficient to generate neurodegenerative effects in females, their combination facilitated the emergence of quantifiable brain deficits. These findings provide an important foundation to further investigate the molecular and cellular mechanisms underlying sex differences in injury responses and neurodegeneration.

Sex-related differences have been documented in many medical conditions, including neurodegenerative disorders ^59-62^ and neurotrauma of various severities ^11,13,63^. Sex differences include anatomical differences, sex gonadal hormones, and immunological responses, all of which presumably affect vulnerability to injury. In fact, there seems to be growing consensus concerning sex-dependent effects in response to TBI, though clinical and pre-clinical findings are inconsistent. While female sex is more often associated with worse outcomes in human TBI studies, in preclinical TBI studies it is more frequently associated with better outcomes and neuroprotection after TBI ^13^. Part of this disparity may be attributed to differences in injury severity and animal model, but the underlying drivers of sex differences in brain injury responses and neurodegenerative conditions remain to be fully elucidated. While sex differences in insects inherently differ from that in humans, there are some similarities associated with reproduction and hormone regulation. Therefore, it is likely that some of the fundamental mechanisms revealed in *Drosophila* may be conserved in humans.

Mating elicits two major types of changes in female flies; increase in egg-laying and reduction in receptivity to mating ^29,52^. Sperm alone can induce a short-term post-mating response, but a sustained response (>7 days) require SP ^29^. SP signaling via SPR activation is believed to switch the female to a state that maximizes reproduction ^53^, which is considered evolutionarily beneficial. However, SP exposure can jeopardize female survival by decreasing daytime sleep ^64,65^, altering the immune system ^66-69^, and increasing female susceptibility to age-dependent tumors ^70^. To our knowledge, the current study is the first to directly show that SP-signaling can elevate vulnerability to injury-induced neurodegeneration in female flies. While mammals do not have SP, reproduction in females is known to dramatically alter metabolism, immune system, hormones, and neurobiology, all of which are vital for maintaining pregnancy. A connection between increased risk for neurodegeneration and hormonal changes associated with reproduction has been suggested in humans and other mammals ^71-73^. There are also contradictory findings, where female sex hormones are found to be neuroprotective ^74,75^. These findings highlight the need for further animal studies with better controls of hormonal and reproductive cycles.

The current study also examines how aging affects response to mild head injury. In humans, age is one of the strongest outcome predictors for complications following head trauma, including mild trauma ^76-78^. Though the cause of injury may differ, older age is consistently associated with an increased incidence of head trauma, a slower overall recovery process, as well as greater morbidity and mortality following injury ^46,48^. Consistent with these observations, we show that flies exposed to vmHT at older ages developed neurodegenerative deficits at a much faster rate than flies injured at younger ages. Our finding that aging accelerates injury-induced neurodegeneration is also supported by findings in humans and other animal models ^79-81^, suggesting conserved pathways underlying aging-induced vulnerability. Previous studies have suggested that mitochondrial function ^82,83^ and immune function ^84,85^ decline with aging. Given that mitochondrial dysfunction ^86,87^ and compromised immune responses ^88,89^ have also been implicated in TBI-induced brain dysfunction and degeneration, future work using our mild injury paradigm should investigate their contributions to aging-increased vulnerability and the development of neurodegeneration in response to head trauma.

Sex differences in aging are conserved across species. In line with this, we demonstrated sex differences in aging-associated vulnerability. While increasing age-at-injury in mated females is generally associated with more severe neurological deficits, females injured on D17 exhibit similar degrees of climbing deficits and vacuolation to those injured on D31, suggesting a possible plateau (Figure 3A). On the other hand, male flies are very resilient to vmHT at younger ages (D3 and D17), displaying little to no phenotypes even on D45 (Figure 3A). However, at the advanced age of 31 days post-eclosion, exposure to vmHT drastically accelerates neurodegeneration in the male brain, resulting in deficits similar to that of mated females of the same injury group (Figure 3B & E). As flies age, male flies seem to show a larger decline in resistance to injury-induced neurodegeneration than females. This finding is supported by work studying sex differences in immune senescence ^90^. In males, this is believed to occur principally due to age-related deterioration in barrier defenses, whereas in females this largely manifests in decreases in innate immunity. Older males may have a weaker exoskeletal defense system, making them more vulnerable to the effects of injury.

To gain some insights into the molecular mechanisms underlying the sexually dimorphic, late-life emergence of neurodegenerative conditions after early exposure to vmHT, we surveyed the transcriptomic landscapes of fly brains with and without vmHT exposure. We were specifically interested in identifying genes whose changes in expression might underlie the observed sex differences between male and mated females, as well as between females of varying reproductive status. Across all conditions and timepoints, we found that the injuries (regardless of age) elicited a very small number of differentially expressed genes that skewed towards decrease in expression. This was initially surprising, because previous *Drosophila* TBI work using the same RNA extraction protocol and sequencing company had reported large numbers of both immediate and lasting gene activation ^91^. When we compared sham males and females at the acute timepoint, we found robust sex-specific gene expression (data not shown) and distinct clustering by sex in our PCA analyses (Figure 6B). This confirmed the validity of our data and subsequent analyses and suggested that the small number of differentially expressed genes was likely due to the mild nature of our injury paradigm. Furthermore, it is possible that not all flies experienced increased neurodegeneration following the mild injuries, which could mask transcriptomic changes due to injury-induced neurodegeneration. Unlike other work utilizing a more severe injury paradigm ^91^, we did not find any acute gene changes that persisted until the very chronic timepoint of 6 weeks following injury, suggesting that mild injury may induce dynamic transcriptomic changes throughout a fly’s lifespan. Additionally, cellular mechanisms that directly contribute to the development of neurodegenerative conditions may not be immediately active following injury; instead, they may be revealed by aging. To better understand the connections and interactions between acute and chronic changes in the transcriptome, future studies may further characterize these dynamic changes at multiple timepoints throughout the fly’s lifespan.

Previous work using a mild injury model has shown that AMP expression is elevated in the brain immediately following injury (in the first 12 hours), returned to baseline at 24 hours post-injury, and then elevated again 1 week post injury ^22^. This suggests that persistent elevation of AMP expression levels and the activation of the innate immune system at a chronic timepoint contributes to neurodegeneration. For both mated females and males, our results comparing injured and sham flies at the acute timepoint (1dpi) do not show any elevation of AMP or AMP-like genes, which is congruent with previous data. A small percentage of immune-related gene changes were identified at this time, but these changes were all biased towards an injury-induced suppression. However, at 6 weeks post injury, we find that the injury paradigm overwhelmingly depressed AMP and AMP-like gene expression in both male and mated female brains but not in virgin and SP-null mated female brains. Interestingly, injury-induced suppression of *Metchnikowin* and *Diptericin* and increased expression of *PPO2* observed at the chronic timepoint is correlated with female-specific late-life behavioral deficits and brain pathology. This finding is supported by a recent report that mated female flies exhibited decreased markers of oxidative stress following trauma, which could be due to immune suppression following a TBI-induced immune challenge ^92^. Additionally, work comparing bacterial infection in virgin, SP^0^-mated, and wildtype-mated females suggest that mating and particularly egg-laying reduces the female’s overall ability to defend against bacterial infections, leading to decreased expression of AMP genes after infections ^68^. This process is mediated by increases in Juvenile Hormone ^66^, which is known to suppress innate immune processes ^93^. SP exposure also acutely activates the Toll-like receptor and immune deficiency (Imd) pathways in females, altering levels of *Metchnikowin* and *Diptericin* ^69^. Furthermore, even one instance of mating is sufficient to induce chronic immunosuppression ^94^. These changes may compound injury-induced changes to disrupt immune homeostasis, leading to observed immunosuppression in mated females. At an advanced age, a decreased expression of AMPs suggest that the mated female fly’s innate defense may be worsened by injury, making them unable to mount a sufficient immune response to pathological insults and therefore more susceptible to infections and disease.

Our finding of the decreased expression of AMP genes at this very chronic timepoint of 6-weeks post-injury, in combination with other groups’ findings of short-term elevation of AMP genes, also suggest a temporally dynamic gene expression pattern in response to head injury. Though immune suppression in mated females seems to contribute to neurodegenerative phenotypes, it is not immediately clear why suppression of a different set of AMP genes was not sufficient for neurodegenerative phenotypes in males. Future work can utilize males injured at older ages or with a more severe paradigm to understand male-specific transcriptome changes that contribute to injury-induced neurodegenerative conditions. Finally, though our RNA-seq data is informative, we acknowledge that there is limited correlation between mRNA and protein levels. Clearly, future studies are needed to dissect the genetic components and molecular players involved in the sex-different development of neurodegenerative conditions after mild head trauma.

## Abbreviations used

HIFLI: <u>H</u>eadfast <u>I</u>mpact <u>FLy</u> <u>I</u>njury
mNGA: <u>m</u>odified <u>N</u>egative <u>G</u>eotaxis <u>A</u>ssay
SP: <u>S</u>ex <u>P</u>eptide
SPR: <u>S</u>ex <u>P</u>eptide <u>R</u>eceptor
SPSN: <u>S</u>ex <u>P</u>eptide <u>S</u>ensory <u>N</u>euron
TBI: <u>T</u>raumatic <u>B</u>rain <u>I</u>njury
vmHT: <u>v</u>ery <u>m</u>ild <u>H</u>ead <u>T</u>rauma

---

**Supplementary Figure S1.**
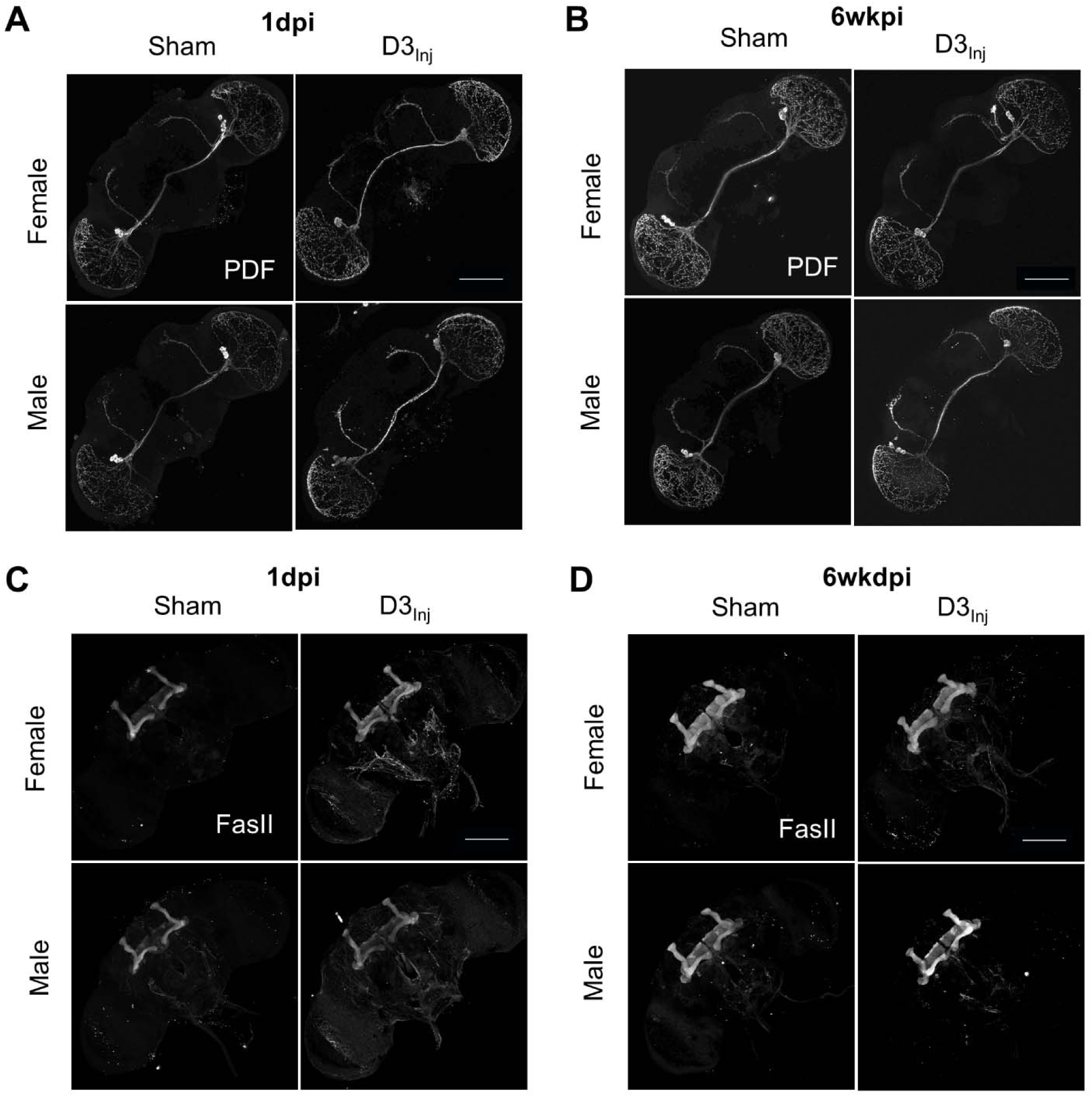
Exposure to vmHT on D3 does not elicit gross morphological alterations in PDF neurons and mushroom bodies. **(A-B)** vmHT exposure on D3 did not significantly alter PDF neuron axonal projections or dendritic arborizations when assessed 24 hours or 6 weeks after injury. **(C-D)** vmHT exposure on D3 did not visibly alter mushroom body axon bundles when assessed 24 hours or 6 weeks after injury. Scale bar = 100 microns.

**Supplementary Figure 2.**
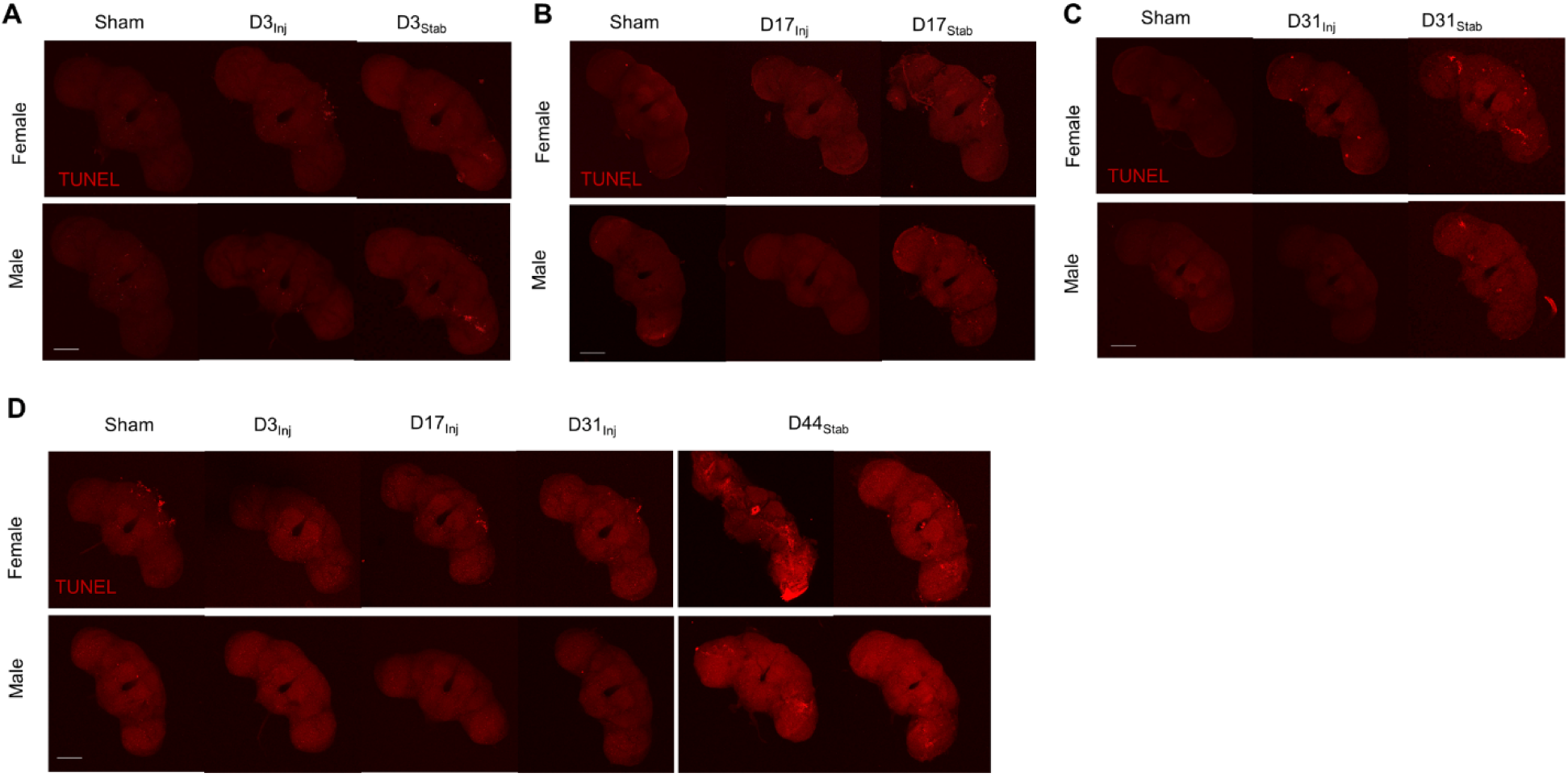
Exposure to vmHT does not elicit significant apoptosis. **(A-C)** Representative images of TUNEL staining in *Drosophila* brains at 1 day post D3_Inj_, D17_Inj_, and D31_Inj_. There was limited TUNEL signal in both sexes and no differences in staining pattern and signal intensity were observed between sham and injured groups. A set of flies were subject to stabbing injuries at the same time and their brains were used as positive controls of apoptosis (right panels). TUNEL staining was visible in the optic lobes or other regions that were penetrated by the fly pin. **(D)** Representative images of TUNEL staining in sham and injured brains on D45. vmHT were delivered on D3, D17, or D31. Stabbing injuries were performed on D44 (24 hours prior to fixation). No differences in staining pattern and signal intensity were observed between sham and injured groups and between sexes. In comparison, brains of stabbed flies (right panels) exhibit much stronger TUNEL signal intensity and region-specific apoptosis near location of primary injury. Scale bar = 100 microns. Each condition contained n=15 flies.

**Supplementary Figure 3.**
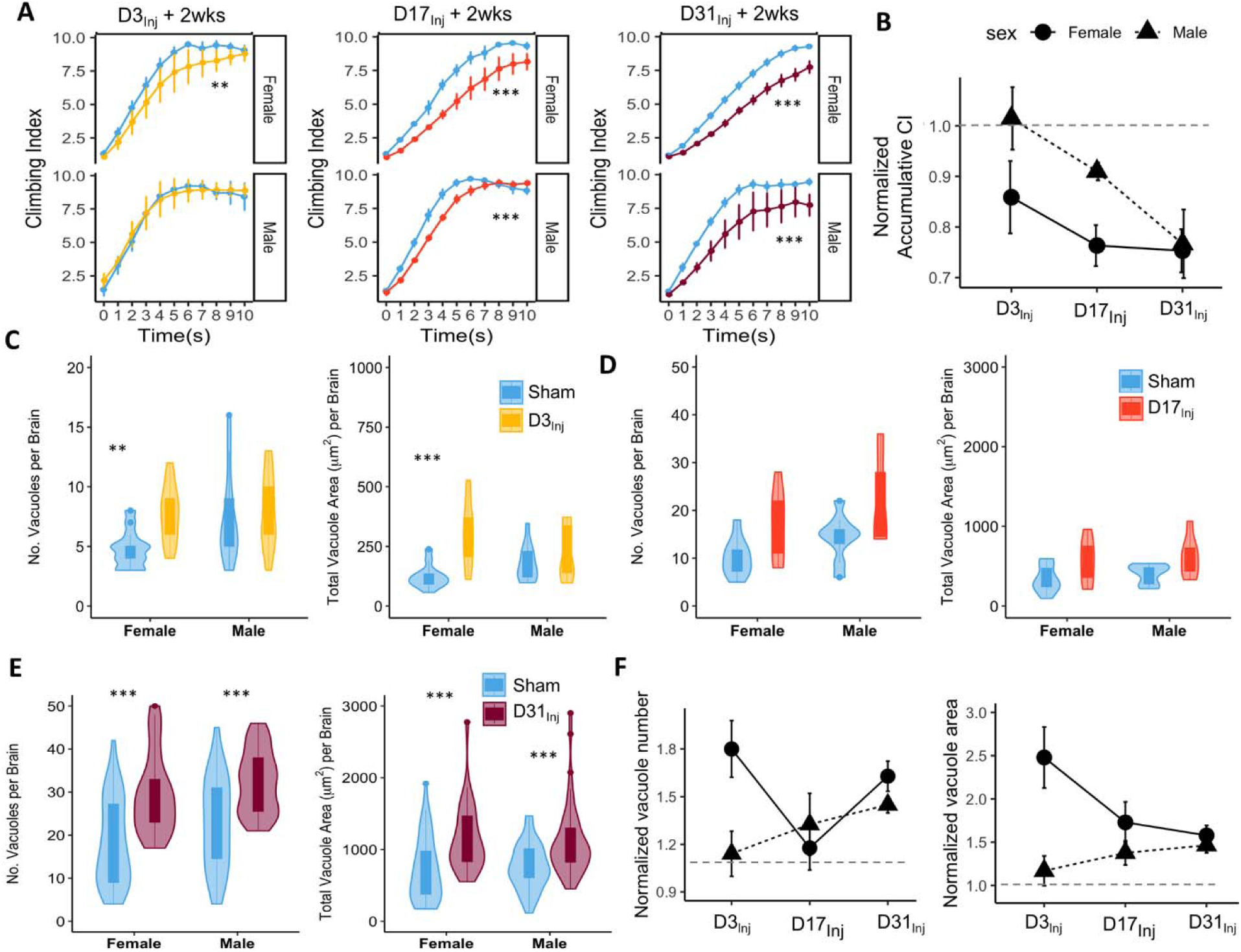
Age-at-injury affects post-injury recovery after two weeks. **(A)** CI plots depicting sensorimotor behavior assessed two weeks following D3_Inj_, D17_Inj_, and D31_Inj_. Female D3_Inj_ groups exhibited a significant decrease in negative geotaxis when assessed on D17 (** p=0.00114), whereas males did not (p>0.05). Both female (*** p= 1.53e-10) and male (*** p= 2.47e-05) D17_Inj_ groups suffered sensorimotor deficits when assessed on D31. D31_Inj_ flies all exhibited significant detriment to climbing behavior when assessed on D45 (*** p= 2e-16 for female, *** p= 2.58e-07 for male). Repeated Measures ANOVA tests were used to calculate statistical significance. Total of three experimental repeats and a total of 9 videos were used in each condition. n>15 per condition per repeat. **(B)** In both male and female flies, older age-at-injury is associated with larger decreases in accumulated climbing indices normalized against the respective sham controls. **(C-E)** Quantification of vacuole formation at two weeks following D3_Inj_, D17_Inj_, and D31_Inj_. D3_Inj_ increased vacuole number (** p=0.0017) and total vacuole area (*** p=0.00016) in female brains but not in male brains. D17_Inj_ did not significantly alter vacuole formation in both sexes though there seems to be a trend towards injury increasing both vacuole size and vacuole number. D31_Inj_ increased both vacuole formation in female and males. Two repeats resulting in n>12 brains per condition. Wilcoxon rank sum tests were used to calculate p values. From left to right, *** p = 2.6e-05, *** p=4.5e-05, *** p=8e-05, and *** p=0.00048. (F) Vacuole number and total vacuole number in each injury condition were normalized against their respective sham groups. Older age-at-injuries were associated with accelerated vacuole formation in males, but this association was not observed in females.

**Supplementary Figure S4.**
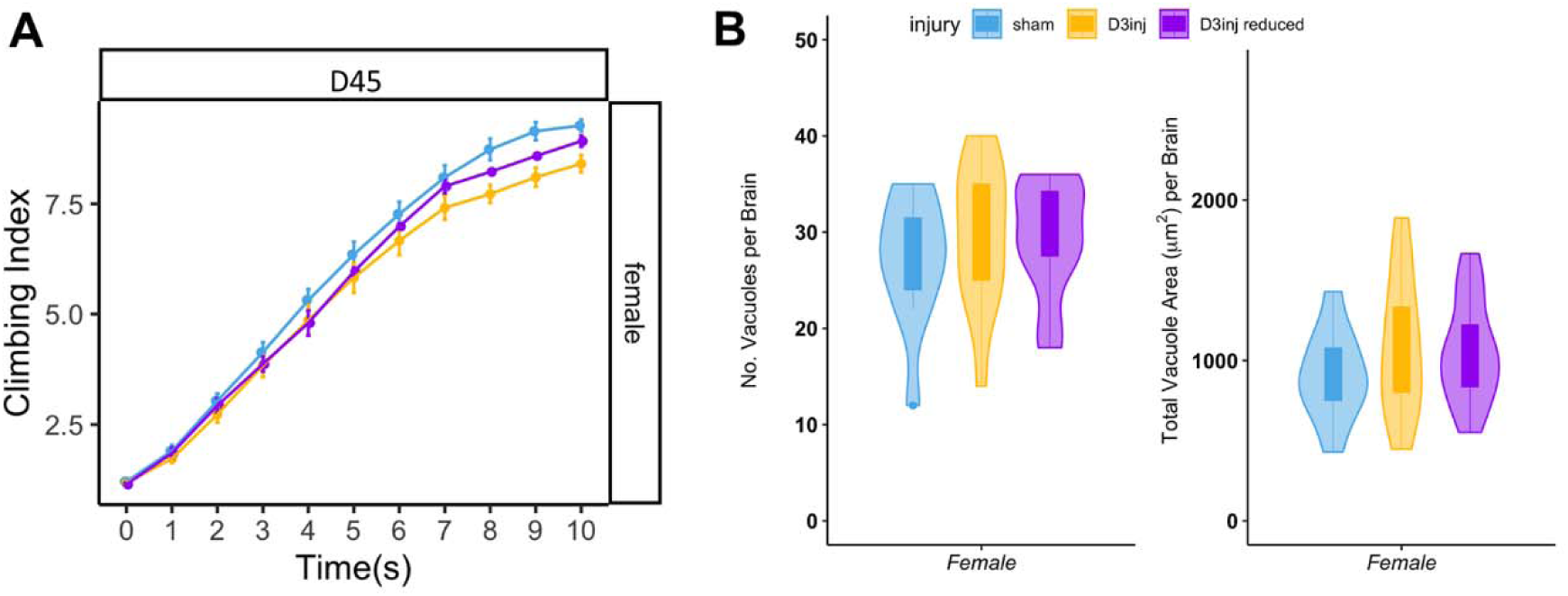
Exposure to reduced vmHT on D3 elicited similar late-life behavioral deficits and brain pathology as exposure to regular vmHT. (A) CI plots depicting decreases in CI in both regular D3_Inj_ and reduced D3_Inj_ females when climbing behavior was assessed on D45. (B) Quantification of vacuole formation on D45. Vacuolation was similarly elevated in reduced D3_Inj_ brains and in regular D3_Inj_ brains. Only one experiment was performed for reduced D3_Inj_ (n=15).

## References

1 Fu, W. Y. & Ip, N. Y. The role of genetic risk factors of Alzheimer’s disease in synaptic dysfunction. Semin Cell Dev Biol (2022). 10.1016/j.semcdb.2022.07.011

2 Pihlstrøm, L., Wiethoff, S. & Houlden, H. Genetics of neurodegenerative diseases: an overview. Handb Clin Neurol 145, 309-323 (2017). 10.1016/b978-0-12-802395-2.00022-5

3 Bertram, L. & Tanzi, R. E. The genetic epidemiology of neurodegenerative disease. J Clin Invest 115, 1449-1457 (2005). 10.1172/jci24761

4 Hardy, J. & Orr, H. The genetics of neurodegenerative diseases. J Neurochem 97, 1690-1699 (2006). 10.1111/j.1471-4159.2006.03979.x

5 Azam, S., Haque, M. E., Balakrishnan, R., Kim, I. S. & Choi, D. K. The Ageing Brain: Molecular and Cellular Basis of Neurodegeneration. Front Cell Dev Biol 9, 683459 (2021). 10.3389/fcell.2021.683459

6 Saikumar, J. & Bonini, N. M. Synergistic effects of brain injury and aging: common mechanisms of proteostatic dysfunction. Trends Neurosci 44, 728-740 (2021). 10.1016/j.tins.2021.06.003

7 Brown, R. C., Lockwood, A. H. & Sonawane, B. R. Neurodegenerative diseases: an overview of environmental risk factors. Environ Health Perspect 113, 1250-1256 (2005). 10.1289/ehp.7567

8 Gupta, R. & Sen, N. Traumatic brain injury: a risk factor for neurodegenerative diseases. Rev Neurosci 27, 93-100 (2016). 10.1515/revneuro-2015-0017

9 Brett, B. L., Gardner, R. C., Godbout, J., Dams-O’Connor, K. & Keene, C. D. Traumatic Brain Injury and Risk of Neurodegenerative Disorder. Biol Psychiatry 91, 498-507 (2022). 10.1016/j.biopsych.2021.05.025

10 Ntikas, M., Binkofski, F., Shah, N. J. & Ietswaart, M. Repeated Sub-Concussive Impacts and the Negative Effects of Contact Sports on Cognition and Brain Integrity. Int J Environ Res Public Health 19 (2022). 10.3390/ijerph19127098

11 Yue, J. K. et al. Age and sex-mediated differences in six-month outcomes after mild traumatic brain injury in young adults: a TRACK-TBI study. Neurological research 41, 609–623 (2019).

12 Berz, K. et al. Sex-specific differences in the severity of symptoms and recovery rate following sports-related concussion in young athletes. The Physician and sportsmedicine 41, 58–63 (2013).

13 Gupte, R., Brooks, W., Vukas, R., Pierce, J. & Harris, J. Sex Differences in Traumatic Brain Injury: What We Know and What We Should Know. Journal of Neurotrauma 36, 3063–3091 (2019). 10.1089/neu.2018.6171

14. 14 Hirth, F. Drosophila melanogaster in the study of human neurodegeneration. CNS & Neurological Disorders-Drug Targets (Formerly Current Drug Targets-CNS & Neurological Disorders) 9, 504-523 (2010).

15 Lessing, D. & Bonini, N. M. Maintaining the brain: insight into human neurodegeneration from Drosophila melanogaster mutants. Nature Reviews Genetics 10, 359–370 (2009).

16 Petersen, A. J. & Wassarman, D. A. Drosophila innate immune response pathways moonlight in neurodegeneration. Fly (Austin) 6, 169-172 (2012). 10.4161/fly.20999

17 McGurk, L., Berson, A. & Bonini, N. M. Drosophila as an In Vivo Model for Human Neurodegenerative Disease. Genetics 201, 377-402 (2015). 10.1534/genetics.115.179457

18 Pandey, U. B. & Nichols, C. D. Human disease models in Drosophila melanogaster and the role of the fly in therapeutic drug discovery. Pharmacol Rev 63, 411-436 (2011). 10.1124/pr.110.003293, 10.1124/pr.110.003293. Epub 2011 Mar 17.

19 Yuva-Aydemir, Y., Almeida, S. & Gao, F. B. Insights into C9ORF72-Related ALS/FTD from Drosophila and iPSC Models. Trends Neurosci 41, 457-469 (2018). 10.1016/j.tins.2018.04.002

20 Bolus, H., Crocker, K., Boekhoff-Falk, G. & Chtarbanova, S. Modeling neurodegenerative disorders in Drosophila melanogaster. International Journal of Molecular Sciences 21, 3055 (2020).

21 Prüßing, K., Voigt, A. & Schulz, J. B. Drosophila melanogaster as a model organism for Alzheimer’s disease. Mol Neurodegener 8, 35 (2013). 10.1186/1750-1326-8-35

22 Barekat, A. et al. Using Drosophila as an integrated model to study mild repetitive traumatic brain injury. Scientific reports 6, 25252 (2016).

23 Katzenberger, R. J. et al. A Drosophila model of closed head traumatic brain injury. Proceedings of the National Academy of Sciences 110, E4152–E4159 (2013).

24 Sun, M. & Chen, L. L. A Novel Method to Model Chronic Traumatic Encephalopathy in Drosophila. Journal of visualized experiments: JoVE (2017).

25 Saikumar, J., Byrns, C. N., Hemphill, M., Meaney, D. F. & Bonini, N. M. Dynamic neural and glial responses of a head-specific model for traumatic brain injury in Drosophila. Proceedings of the National Academy of Sciences 117, 17269–17277 (2020).

26 van Alphen, B. et al. Glial immune-related pathways mediate effects of closed head traumatic brain injury on behavior and lethality in Drosophila. PLoS Biol 20, e3001456 (2022). 10.1371/journal.pbio.3001456

27 Ye, C., Behnke, J. A., Hardin, K. R. & Zheng, J. Q. Drosophila melanogaster as a model to study age and sex differences in brain injury and neurodegeneration after mild head trauma. Frontiers in Neuroscience 17 (2023). 10.3389/fnins.2023.1150694

28 Behnke, J. A., Ye, C., Setty, A., Moberg, K. H. & Zheng, J. Q. Repetitive mild head trauma induces activity mediated lifelong brain deficits in a novel Drosophila model. Scientific Reports 11, 9738 (2021). 10.1038/s41598-021-89121-7

29 Liu, H. & Kubli, E. Sex-peptide is the molecular basis of the sperm effect in Drosophila melanogaster. Proc Natl Acad Sci U S A 100, 9929-9933 (2003). 10.1073/pnas.1631700100

30 Wang, F. et al. Neural circuitry linking mating and egg laying in Drosophila females. Nature 579, 101-105 (2020). 10.1038/s41586-020-2055-9

31 Romero-Ferrero, F., Bergomi, M. G., Hinz, R. C., Heras, F. J. H. & de Polavieja, G. G. idtracker.ai: tracking all individuals in small or large collectives of unmarked animals. Nature Methods 16, 179-182 (2019). 10.1038/s41592-018-0295-5

32 Behnke, J. A., Ye, C., Moberg, K. H. & Zheng, J. Q. A protocol to detect neurodegeneration in Drosophila melanogaster whole-brain mounts using advanced microscopy. STAR Protoc 2, 100689 (2021). 10.1016/j.xpro.2021.100689

33 Afgan, E. et al. The Galaxy platform for accessible, reproducible and collaborative biomedical analyses: 2018 update. Nucleic Acids Res 46, W537-w544 (2018). 10.1093/nar/gky379

34 Andrews, S. FastQC: a quality control tool for high throughput sequence data. Available online at: http://www.bioinformatics.babraham.ac.uk/projects/fastqc, 2010).

35 Dobin, A. et al. STAR: ultrafast universal RNA-seq aligner. Bioinformatics 29, 15-21 (2013). 10.1093/bioinformatics/bts635

36 Cunningham, F. et al. Ensembl 2022. Nucleic Acids Research 50, D988–D995 (2021). 10.1093/nar/gkab1049

37 Robinson, J. T. et al. Integrative genomics viewer. Nat Biotechnol 29, 24–26 (2011). 10.1038/nbt.1754

38 Danecek, P. et al. Twelve years of SAMtools and BCFtools. GigaScience 10 (2021). 10.1093/gigascience/giab008

39 Wang, L., Wang, S. & Li, W. RSeQC: quality control of RNA-seq experiments. Bioinformatics 28, 2184-2185 (2012). 10.1093/bioinformatics/bts356

40 Liao, Y., Smyth, G. K. & Shi, W. featureCounts: an efficient general purpose program for assigning sequence reads to genomic features. Bioinformatics 30, 923–930 (2014). 10.1093/bioinformatics/btt656

41 Love, M. I., Huber, W. & Anders, S. Moderated estimation of fold change and dispersion for RNA-seq data with DESeq2. Genome Biology 15, 550 (2014). 10.1186/s13059-014-0550-8

42 Wu, T. et al. clusterProfiler 4.0: A universal enrichment tool for interpreting omics data. The Innovation 2 (2021). 10.1016/j.xinn.2021.100141

43 Gramates, L. S. et al. FlyBase: a guided tour of highlighted features. Genetics 220 (2022). 10.1093/genetics/iyac035

44 Katzenberger, R. J., Ganetzky, B. & Wassarman, D. A. Age and Diet Affect Genetically Separable Secondary Injuries that Cause Acute Mortality Following Traumatic Brain Injury in Drosophila. G3 (Bethesda) 6, 4151–4166 (2016). 10.1534/g3.116.036194

45 Katzenberger, R. J. et al. A Drosophila model of closed head traumatic brain injury. Proc Natl Acad Sci U S A 110, E4152–4159 (2013). 10.1073/pnas.1316895110

46 Abdulle, A. E. et al. Early predictors for long-term functional outcome after mild traumatic brain injury in frail elderly patients. The Journal of Head Trauma Rehabilitation 33, E59-E67 (2018).

47 Peterson, A. B., Xu, L., Daugherty, J. & Breiding, M. J. Surveillance report of traumatic brain injury-related emergency department visits, hospitalizations, and deaths, United States, 2014. (2019).

48 Mosenthal, A. C. et al. The effect of age on functional outcome in mild traumatic brain injury: 6-month report of a prospective multicenter trial. Journal of Trauma and Acute Care Surgery 56, 1042–1048 (2004).

49 Heisenberg, M. & Böhl, K. Isolation of anatomical brain mutants of Drosophila by histological means. Zeitschrift für Naturforschung C 34, 143–147 (1979).

50 Sunderhaus, E. R. & Kretzschmar, D. Mass histology to quantify neurodegeneration in Drosophila. JoVE (Journal of Visualized Experiments), e54809 (2016).

51 Sanuki, R. Drosophila models of traumatic brain injury. FBL 25, 168–178 (2020). 10.2741/4801

52 Chapman, T. et al. The sex peptide of Drosophila melanogaster: female post-mating responses analyzed by using RNA interference. Proc Natl Acad Sci U S A 100, 9923–9928 (2003). 10.1073/pnas.1631635100

53 Yapici, N., Kim, Y. J., Ribeiro, C. & Dickson, B. J. A receptor that mediates the post-mating switch in Drosophila reproductive behaviour. Nature 451, 33–37 (2008). 10.1038/nature06483

54 Hasemeyer, M., Yapici, N., Heberlein, U. & Dickson, B. J. Sensory neurons in the Drosophila genital tract regulate female reproductive behavior. Neuron 61, 511–518 (2009). 10.1016/j.neuron.2009.01.009

55 Rezaval, C. et al. Neural circuitry underlying Drosophila female postmating behavioral responses. Curr Biol 22, 1155–1165 (2012). 10.1016/j.cub.2012.04.062

56 Yang, C. H. et al. Control of the postmating behavioral switch in Drosophila females by internal sensory neurons. Neuron 61, 519-526 (2009). 10.1016/j.neuron.2008.12.021

57 Hanson, M. A. & Lemaitre, B. Antimicrobial peptides do not directly contribute to aging in Drosophila, but improve lifespan by preventing dysbiosis. Dis Model Mech 16 (2023). 10.1242/dmm.049965

58 Binggeli, O., Neyen, C., Poidevin, M. & Lemaitre, B. Prophenoloxidase Activation Is Required for Survival to Microbial Infections in Drosophila. PLOS Pathogens 10, e1004067 (2014). 10.1371/journal.ppat.1004067

59 Vegeto, E. et al. The Role of Sex and Sex Hormones in Neurodegenerative Diseases. Endocr Rev 41, 273–319 (2020). 10.1210/endrev/bnz005

60 Cerri, S., Mus, L. & Blandini, F. Parkinson’s Disease in Women and Men: What’s the Difference? J Parkinsons Dis 9, 501-515 (2019). 10.3233/jpd-191683

61 Nebel, R. A. et al. Understanding the impact of sex and gender in Alzheimer’s disease: A call to action. Alzheimers Dement 14, 1171–1183 (2018). 10.1016/j.jalz.2018.04.008

62 Podcasy, J. L. & Epperson, C. N. Considering sex and gender in Alzheimer disease and other dementias. Dialogues Clin Neurosci 18, 437–446 (2016). 10.31887/DCNS.2016.18.4/cepperson

63 Rauen, K., et al. Quality of life after traumatic brain injury: a cross-sectional analysis uncovers age-and sex-related differences over the adult life span. GeroScience, 1–16 (2020).

64 Isaac, R. E., Li, C., Leedale, A. E. & Shirras, A. D. Drosophila male sex peptide inhibits siesta sleep and promotes locomotor activity in the post-mated female. Proc Biol Sci 277, 65–70 (2010). 10.1098/rspb.2009.1236

65 Garbe, D. S. et al. Changes in Female Drosophila Sleep following Mating Are Mediated by SPSN-SAG Neurons. J Biol Rhythms 31, 551–567 (2016). 10.1177/0748730416668048

66 Schwenke, R. A. & Lazzaro, B. P. Juvenile Hormone Suppresses Resistance to Infection in Mated Female Drosophila melanogaster. Curr Biol 27, 596–601 (2017). 10.1016/j.cub.2017.01.004

67 Short, S. M. & Lazzaro, B. P. Female and male genetic contributions to post-mating immune defence in female Drosophila melanogaster. Proc Biol Sci 277, 3649–3657 (2010). 10.1098/rspb.2010.0937

68 Short, S. M., Wolfner, M. F. & Lazzaro, B. P. Female Drosophila melanogaster suffer reduced defense against infection due to seminal fluid components. J Insect Physiol 58, 1192–1201 (2012). 10.1016/j.jinsphys.2012.06.002

69 Peng, J., Zipperlen, P. & Kubli, E. Drosophila sex-peptide stimulates female innate immune system after mating via the Toll and Imd pathways. Curr Biol 15, 1690–1694 (2005). 10.1016/j.cub.2005.08.048

70 Regan, J. C. et al. Sex difference in pathology of the ageing gut mediates the greater response of female lifespan to dietary restriction. Elife 5, e10956 (2016). 10.7554/eLife.10956

71 Xiong, J. et al. FSH blockade improves cognition in mice with Alzheimer’s disease. Nature 603, 470–476 (2022). 10.1038/s41586-022-04463-0

72 Gilsanz, P. et al. Reproductive period and risk of dementia in a diverse cohort of health care members. Neurology 92, e2005–e2014 (2019). 10.1212/wnl.0000000000007326

73 Jang, H. et al. Differential effects of completed and incomplete pregnancies on the risk of Alzheimer disease. Neurology 91, e643–e651 (2018). 10.1212/wnl.0000000000006000

74 Ha, D. M., Xu, J. & Janowsky, J. S. Preliminary evidence that long-term estrogen use reduces white matter loss in aging. Neurobiol Aging 28, 1936–1940 (2007). 10.1016/j.neurobiolaging.2006.08.007

75 Song, Y. J. et al. The Effect of Estrogen Replacement Therapy on Alzheimer’s Disease and Parkinson’s Disease in Postmenopausal Women: A Meta-Analysis. Front Neurosci 14, 157 (2020). 10.3389/fnins.2020.00157

76 Moretti, L. et al. Cognitive decline in older adults with a history of traumatic brain injury. The Lancet Neurology 11, 1103–1112 (2012).

77 Jacobs, B. et al. Outcome prediction in mild traumatic brain injury: age and clinical variables are stronger predictors than CT abnormalities. Journal of neurotrauma 27, 655–668 (2010).

78 Bittencourt- Villalpando, M., van der Horn, H. J., Maurits, N. M. & van der Naalt, J. Disentangling the effects of age and mild traumatic brain injury on brain network connectivity: A resting state fMRI study. Neuroimage Clin 29, 102534 (2020). 10.1016/j.nicl.2020.102534

79 Doust, Y. V., Rowe, R. K., Adelson, P. D., Lifshitz, J. & Ziebell, J. M. Age-at-Injury Determines the Extent of Long-Term Neuropathology and Microgliosis After a Diffuse Brain Injury in Male Rats. Frontiers in Neurology 12 (2021). 10.3389/fneur.2021.722526

80 Rowe, R. K. et al. Aging with Traumatic Brain Injury: Effects of Age at Injury on Behavioral Outcome following Diffuse Brain Injury in Rats. Dev Neurosci 38, 195-205 (2016). 10.1159/000446773

81 Ojo, J. O. et al. Repetitive mild traumatic brain injury augments tau pathology and glial activation in aged hTau mice. J Neuropathol Exp Neurol 72, 137-151 (2013). 10.1097/NEN.0b013e3182814cdf

82 Cui, H., Kong, Y. & Zhang, H. Oxidative stress, mitochondrial dysfunction, and aging. Journal of signal transduction 2012 (2012).

83 Chistiakov, D. A., Sobenin, I. A., Revin, V. V., Orekhov, A. N. & Bobryshev, Y. V. Mitochondrial aging and age-related dysfunction of mitochondria. BioMed research international 2014 (2014).

84 Weyand, C. M. & Goronzy, J. J. Aging of the Immune System. Mechanisms and Therapeutic Targets. Ann Am Thorac Soc 13 **Suppl 5**, S422-s428 (2016). 10.1513/AnnalsATS.201602-095AW

85 Haynes, L. Aging of the Immune System: Research Challenges to Enhance the Health Span of Older Adults. Frontiers in Aging 1 (2020). 10.3389/fragi.2020.602108

86 Cheng, G., Kong, R. H., Zhang, L. M. & Zhang, J. N. Mitochondria in traumatic brain injury and mitochondrial-targeted multipotential therapeutic strategies. Br J Pharmacol 167, 699-719 (2012). 10.1111/j.1476-5381.2012.02025.x

87 Kim, S., Han, S. C., Gallan, A. J. & Hayes, J. P. Neurometabolic indicators of mitochondrial dysfunction in repetitive mild traumatic brain injury. Concussion 2, CNC45 (2017). 10.2217/cnc-2017-0013

88 Needham, E. J. et al. The immunological response to traumatic brain injury. J Neuroimmunol 332, 112-125 (2019). 10.1016/j.jneuroim.2019.04.005

89 Verboon, L. N., Patel, H. C. & Greenhalgh, A. D. The Immune System’s Role in the Consequences of Mild Traumatic Brain Injury (Concussion). Frontiers in Immunology 12 (2021). 10.3389/fimmu.2021.620698

90 Kubiak, M. & Tinsley, M. C. Sex-Specific Routes To Immune Senescence In Drosophila melanogaster. Sci Rep 7, 10417 (2017). 10.1038/s41598-017-11021-6

91 Byrns, C. N., Saikumar, J. & Bonini, N. M. Glial AP1 is activated with aging and accelerated by traumatic brain injury. Nat Aging 1, 585-597 (2021). 10.1038/s43587-021-00072-0

92 Jones, T. B. et al. Mild traumatic brain injury in Drosophila melanogaster alters reactive oxygen and nitrogen species in a sex-dependent manner. Experimental Neurology 372, 114621 (2024). 10.1016/j.expneurol.2023.114621

93 Flatt, T. et al. Hormonal regulation of the humoral innate immune response in Drosophila melanogaster. J Exp Biol 211, 2712-2724 (2008). 10.1242/jeb.014878

94 Gordon, K. E., Wolfner, M. F. & Lazzaro, B. P. A single mating is sufficient to induce persistent reduction of immune defense in mated female Drosophila melanogaster. J Insect Physiol 140, 104414 (2022). 10.1016/j.jinsphys.2022.104414

