## Supplemental datasets1-3 for "Sexual Dimorphism in Age-Dependent Neurodegeneration After Mild Head Trauma in *Drosophila*: Unveiling the Adverse Impact of Female Reproductive Signaling"

**Supplementary Data S1**

### 1. Statistical comparisons for mNGA on D45 using repeated measures ANOVA

#### A. Female flies, comparing sham, D3, D17, and D31Inj.

```
Error: factor(Time)
          Df Sum Sq Mean Sq F value Pr(>F)
Residuals 10   2154   215.4

Error: Within
          Df Sum Sq Mean Sq F value Pr(>F)
factor(injury)   3  120.0   40.01   62.56 <2e-16 ***
Residuals      327  209.1    0.64
Signif. codes:  0 '***' 0.001 '**' 0.01 '*' 0.05 '.' 0.1 ' ' 1
```

Pair-wise comparisons:

| Time | .y. | group1 | group2 | n1 | n2 | p | p.signif | p.adj | p.adj.signif |
|---|---|---|---|---|---|---|---|---|---|
| 0s | CI | Sham | D3Inj | 9 | 9 | 0.74 | ns | 1 | ns |
| 0s | CI | Sham | D17Inj | 9 | 9 | 0.0803 | ns | 0.482 | ns |
| 0s | CI | D3Inj | D17Inj | 9 | 9 | 0.124 | ns | 0.742 | ns |
| 0s | CI | Sham | D31Inj | 9 | 9 | 0.257 | ns | 1 | ns |
| 0s | CI | D3Inj | D31Inj | 9 | 9 | 0.408 | ns | 1 | ns |
| 0s | CI | D17Inj | D31Inj | 9 | 9 | 0.348 | ns | 1 | ns |
| 1s | CI | Sham | D3Inj | 9 | 9 | 0.247 | ns | 1 | ns |
| 1s | CI | Sham | D17Inj | 9 | 9 | 0.0293 | * | 0.176 | ns |
| 1s | CI | D3Inj | D17Inj | 9 | 9 | 0.146 | ns | 0.874 | ns |
| 1s | CI | Sham | D31Inj | 9 | 9 | 0.0031 | ** | 0.0186 | * |
| 1s | CI | D3Inj | D31Inj | 9 | 9 | 0.0421 | * | 0.253 | ns |
| 1s | CI | D17Inj | D31Inj | 9 | 9 | 0.97 | ns | 1 | ns |
| 2s | CI | Sham | D3Inj | 9 | 9 | 0.171 | ns | 1 | ns |
| 2s | CI | Sham | D17Inj | 9 | 9 | 0.0109 | * | 0.0655 | ns |
| 2s | CI | D3Inj | D17Inj | 9 | 9 | 0.0865 | ns | 0.519 | ns |
| 2s | CI | Sham | D31Inj | 9 | 9 | 0.000263 | *** | 0.00158 | ** |
| 2s | CI | D3Inj | D31Inj | 9 | 9 | 0.00786 | ** | 0.0471 | * |
| 2s | CI | D17Inj | D31Inj | 9 | 9 | 0.78 | ns | 1 | ns |
| 3s | CI | Sham | D3Inj | 9 | 9 | 0.335 | ns | 1 | ns |
| 3s | CI | Sham | D17Inj | 9 | 9 | 0.0268 | * | 0.161 | ns |
| 3s | CI | D3Inj | D17Inj | 9 | 9 | 0.106 | ns | 0.634 | ns |
| 3s | CI | Sham | D31Inj | 9 | 9 | 0.000286 | *** | 0.00172 | ** |
| 3s | CI | D3Inj | D31Inj | 9 | 9 | 0.00318 | ** | 0.0191 | * |
| 3s | CI | D17Inj | D31Inj | 9 | 9 | 0.527 | ns | 1 | ns |
| 4s | CI | Sham | D3Inj | 9 | 9 | 0.263 | ns | 1 | ns |

| | | | | | | | | | |
|---|---|---|---|---|---|---|---|---|---|
| 4s | CI | Sham | D17Inj | 9 | 9 | 0.0196 | * | 0.118 | ns |
| 4s | CI | D3Inj | D17Inj | 9 | 9 | 0.0998 | ns | 0.599 | ns |
| 4s | CI | Sham | D31Inj | 9 | 9 | 0.000297 | *** | 0.00178 | ** |
| 4s | CI | D3Inj | D31Inj | 9 | 9 | 0.00481 | ** | 0.0288 | * |
| 4s | CI | D17Inj | D31Inj | 9 | 9 | 0.624 | ns | 1 | ns |
| 5s | CI | Sham | D3Inj | 9 | 9 | 0.206 | ns | 1 | ns |
| 5s | CI | Sham | D17Inj | 9 | 9 | 0.0069 | ** | 0.0414 | * |
| 5s | CI | D3Inj | D17Inj | 9 | 9 | 0.0508 | ns | 0.305 | ns |
| 5s | CI | Sham | D31Inj | 9 | 9 | 0.000234 | *** | 0.0014 | ** |
| 5s | CI | D3Inj | D31Inj | 9 | 9 | 0.00551 | ** | 0.033 | * |
| 5s | CI | D17Inj | D31Inj | 9 | 9 | 0.9 | ns | 1 | ns |
| 6s | CI | Sham | D3Inj | 9 | 9 | 0.167 | ns | 1 | ns |
| 6s | CI | Sham | D17Inj | 9 | 9 | 0.00354 | ** | 0.0212 | * |
| 6s | CI | D3Inj | D17Inj | 9 | 9 | 0.0341 | * | 0.205 | ns |
| 6s | CI | Sham | D31Inj | 9 | 9 | 0.000177 | *** | 0.00106 | ** |
| 6s | CI | D3Inj | D31Inj | 9 | 9 | 0.00565 | ** | 0.0339 | * |
| 6s | CI | D17Inj | D31Inj | 9 | 9 | 0.95 | ns | 1 | ns |
| 7s | CI | Sham | D3Inj | 9 | 9 | 0.118 | ns | 0.707 | ns |
| 7s | CI | Sham | D17Inj | 9 | 9 | 0.00383 | ** | 0.023 | * |
| 7s | CI | D3Inj | D17Inj | 9 | 9 | 0.0485 | * | 0.291 | ns |
| 7s | CI | Sham | D31Inj | 9 | 9 | 0.00019 | *** | 0.00114 | ** |
| 7s | CI | D3Inj | D31Inj | 9 | 9 | 0.00942 | ** | 0.0565 | ns |
| 7s | CI | D17Inj | D31Inj | 9 | 9 | 0.96 | ns | 1 | ns |
| 8s | CI | Sham | D3Inj | 9 | 9 | 0.0253 | * | 0.152 | ns |
| 8s | CI | Sham | D17Inj | 9 | 9 | 0.00745 | ** | 0.0447 | * |
| 8s | CI | D3Inj | D17Inj | 9 | 9 | 0.21 | ns | 1 | ns |
| 8s | CI | Sham | D31Inj | 9 | 9 | 0.000153 | *** | 0.00092 | *** |
| 8s | CI | D3Inj | D31Inj | 9 | 9 | 0.0394 | * | 0.236 | ns |
| 8s | CI | D17Inj | D31Inj | 9 | 9 | 0.791 | ns | 1 | ns |
| 9s | CI | Sham | D3Inj | 9 | 9 | 0.0182 | * | 0.109 | ns |
| 9s | CI | Sham | D17Inj | 9 | 9 | 0.0087 | ** | 0.0522 | ns |
| 9s | CI | D3Inj | D17Inj | 9 | 9 | 0.272 | ns | 1 | ns |
| 9s | CI | Sham | D31Inj | 9 | 9 | 0.000121 | *** | 0.000725 | *** |
| 9s | CI | D3Inj | D31Inj | 9 | 9 | 0.0437 | * | 0.262 | ns |
| 9s | CI | D17Inj | D31Inj | 9 | 9 | 0.698 | ns | 1 | ns |
| 10s | CI | Sham | D3Inj | 9 | 9 | 0.0356 | * | 0.213 | ns |
| 10s | CI | Sham | D17Inj | 9 | 9 | 0.0114 | * | 0.0686 | ns |
| 10s | CI | D3Inj | D17Inj | 9 | 9 | 0.236 | ns | 1 | ns |
| 10s | CI | Sham | D31Inj | 9 | 9 | 0.00101 | ** | 0.00608 | ** |
| 10s | CI | D3Inj | D31Inj | 9 | 9 | 0.122 | ns | 0.73 | ns |
| 10s | CI | D17Inj | D31Inj | 9 | 9 | 0.954 | ns | 1 | ns |

**B. Male flies, comparing sham, D3, D17, and D31Inj.**

```
Error: factor(Time)
          Df Sum Sq Mean Sq F value Pr(>F)
Residuals 10  934.4   93.44

Error: Within
          Df Sum Sq Mean Sq F value  Pr(>F)
factor(injury)   3  57.22  19.074   22.64 1.2e-11 ***
Residuals      118  99.43   0.843
Signif. codes:  0 ‘***’ 0.001 ‘**’ 0.01 ‘*’ 0.05 ‘.’ 0.1 ‘ ’ 1
```

| Time | .y. | group1 | group2 | n1 | n2 | p | p.signif | p.adj | p.adj.signif |
|---|---|---|---|---|---|---|---|---|---|
| 0s | CI | Sham | D3Inj | 9 | 9 | 0.761 | ns | 1 | ns |
| 0s | CI | Sham | D17Inj | 9 | 9 | 0.122 | ns | 0.735 | ns |
| 0s | CI | D3Inj | D17Inj | 9 | 9 | 0.0755 | ns | 0.453 | ns |
| 0s | CI | Sham | D31Inj | 9 | 9 | 0.15 | ns | 0.901 | ns |
| 0s | CI | D3Inj | D31Inj | 9 | 9 | 0.093 | ns | 0.558 | ns |
| 0s | CI | D17Inj | D31Inj | 9 | 9 | 0.896 | ns | 1 | ns |
| 1s | CI | Sham | D3Inj | 9 | 9 | 0.389 | ns | 1 | ns |
| 1s | CI | Sham | D17Inj | 9 | 9 | 0.00604 | ** | 0.0363 | * |
| 1s | CI | D3Inj | D17Inj | 9 | 9 | 0.0236 | * | 0.141 | ns |
| 1s | CI | Sham | D31Inj | 9 | 9 | 0.005 | ** | 0.03 | * |
| 1s | CI | D3Inj | D31Inj | 9 | 9 | 0.0192 | * | 0.115 | ns |
| 1s | CI | D17Inj | D31Inj | 9 | 9 | 0.897 | ns | 1 | ns |
| 2s | CI | Sham | D3Inj | 9 | 9 | 0.428 | ns | 1 | ns |
| 2s | CI | Sham | D17Inj | 9 | 9 | 0.00793 | ** | 0.0476 | * |
| 2s | CI | D3Inj | D17Inj | 9 | 9 | 0.028 | * | 0.168 | ns |
| 2s | CI | Sham | D31Inj | 9 | 9 | 0.00166 | ** | 0.00999 | ** |
| 2s | CI | D3Inj | D31Inj | 9 | 9 | 0.0052 | ** | 0.0312 | * |
| 2s | CI | D17Inj | D31Inj | 9 | 9 | 0.292 | ns | 1 | ns |
| 3s | CI | Sham | D3Inj | 9 | 9 | 0.789 | ns | 1 | ns |
| 3s | CI | Sham | D17Inj | 9 | 9 | 0.0834 | ns | 0.5 | ns |
| 3s | CI | D3Inj | D17Inj | 9 | 9 | 0.127 | ns | 0.765 | ns |
| 3s | CI | Sham | D31Inj | 9 | 9 | 0.0168 | * | 0.101 | ns |
| 3s | CI | D3Inj | D31Inj | 9 | 9 | 0.0257 | * | 0.154 | ns |
| 3s | CI | D17Inj | D31Inj | 9 | 9 | 0.331 | ns | 1 | ns |
| 4s | CI | Sham | D3Inj | 9 | 9 | 0.778 | ns | 1 | ns |
| 4s | CI | Sham | D17Inj | 9 | 9 | 0.188 | ns | 1 | ns |
| 4s | CI | D3Inj | D17Inj | 9 | 9 | 0.285 | ns | 1 | ns |
| 4s | CI | Sham | D31Inj | 9 | 9 | 0.0387 | * | 0.232 | ns |
| 4s | CI | D3Inj | D31Inj | 9 | 9 | 0.061 | ns | 0.366 | ns |
| 4s | CI | D17Inj | D31Inj | 9 | 9 | 0.332 | ns | 1 | ns |

| | | | | | | | | | |
|---|---|---|---|---|---|---|---|---|---|
| 5s | CI | Sham | D3Inj | 9 | 9 | 0.724 | ns | 1 | ns |
| 5s | CI | Sham | D17Inj | 9 | 9 | 0.373 | ns | 1 | ns |
| 5s | CI | D3Inj | D17Inj | 9 | 9 | 0.58 | ns | 1 | ns |
| 5s | CI | Sham | D31Inj | 9 | 9 | 0.0651 | ns | 0.39 | ns |
| 5s | CI | D3Inj | D31Inj | 9 | 9 | 0.114 | ns | 0.687 | ns |
| 5s | CI | D17Inj | D31Inj | 9 | 9 | 0.267 | ns | 1 | ns |
| 6s | CI | Sham | D3Inj | 9 | 9 | 0.77 | ns | 1 | ns |
| 6s | CI | Sham | D17Inj | 9 | 9 | 0.493 | ns | 1 | ns |
| 6s | CI | D3Inj | D17Inj | 9 | 9 | 0.688 | ns | 1 | ns |
| 6s | CI | Sham | D31Inj | 9 | 9 | 0.108 | ns | 0.648 | ns |
| 6s | CI | D3Inj | D31Inj | 9 | 9 | 0.17 | ns | 1 | ns |
| 6s | CI | D17Inj | D31Inj | 9 | 9 | 0.307 | ns | 1 | ns |
| 7s | CI | Sham | D3Inj | 9 | 9 | 0.951 | ns | 1 | ns |
| 7s | CI | Sham | D17Inj | 9 | 9 | 0.862 | ns | 1 | ns |
| 7s | CI | D3Inj | D17Inj | 9 | 9 | 0.814 | ns | 1 | ns |
| 7s | CI | Sham | D31Inj | 9 | 9 | 0.121 | ns | 0.724 | ns |
| 7s | CI | D3Inj | D31Inj | 9 | 9 | 0.11 | ns | 0.657 | ns |
| 7s | CI | D17Inj | D31Inj | 9 | 9 | 0.158 | ns | 0.949 | ns |
| 8s | CI | Sham | D3Inj | 9 | 9 | 0.918 | ns | 1 | ns |
| 8s | CI | Sham | D17Inj | 9 | 9 | 0.689 | ns | 1 | ns |
| 8s | CI | D3Inj | D17Inj | 9 | 9 | 0.616 | ns | 1 | ns |
| 8s | CI | Sham | D31Inj | 9 | 9 | 0.133 | ns | 0.795 | ns |
| 8s | CI | D3Inj | D31Inj | 9 | 9 | 0.113 | ns | 0.677 | ns |
| 8s | CI | D17Inj | D31Inj | 9 | 9 | 0.244 | ns | 1 | ns |
| 9s | CI | Sham | D3Inj | 9 | 9 | 0.882 | ns | 1 | ns |
| 9s | CI | Sham | D17Inj | 9 | 9 | 0.61 | ns | 1 | ns |
| 9s | CI | D3Inj | D17Inj | 9 | 9 | 0.513 | ns | 1 | ns |
| 9s | CI | Sham | D31Inj | 9 | 9 | 0.166 | ns | 0.994 | ns |
| 9s | CI | D3Inj | D31Inj | 9 | 9 | 0.132 | ns | 0.79 | ns |
| 9s | CI | D17Inj | D31Inj | 9 | 9 | 0.349 | ns | 1 | ns |
| 10s | CI | Sham | D3Inj | 9 | 9 | 0.624 | ns | 1 | ns |
| 10s | CI | Sham | D17Inj | 9 | 9 | 0.546 | ns | 1 | ns |
| 10s | CI | D3Inj | D17Inj | 9 | 9 | 0.907 | ns | 1 | ns |
| 10s | CI | Sham | D31Inj | 9 | 9 | 0.0515 | ns | 0.309 | ns |
| 10s | CI | D3Inj | D31Inj | 9 | 9 | 0.113 | ns | 0.68 | ns |
| 10s | CI | D17Inj | D31Inj | 9 | 9 | 0.136 | ns | 0.817 | ns |

**2. Statistical analyses of vacuole formation using Wilcoxon rank sum tests.**

**A. Comparing number of vacuoles in female brains of sham, D3Inj, D17Inj, and D31Inj conditions.**

Kruskal-Wallis rank sum test

data: total.no by injury
Kruskal-Wallis chi-squared = 26.285, df = 3, p-value = 8.312e-06

Pairwise comparisons using Wilcoxon rank sum test with continuity correction

data: data.F$total.no and data.F$injury

| | sham | D3inj | D17inj |
|---|---|---|---|
| D3inj | 0.01165 | - | - |
| D17inj | 0.00029 | 0.21596 | - |
| D31inj | 0.00015 | 0.21596 | 0.81813 |

**B. Comparing total size of vacuoles in female brains of sham, D3Inj, D17Inj, and D31Inj conditions.**

Kruskal-Wallis rank sum test

data: total.area by injury
Kruskal-Wallis chi-squared = 23.506, df = 3, p-value = 3.167e-05

Pairwise comparisons using Wilcoxon rank sum exact test

data: data.F$total.area and data.F$injury

| | sham | D3inj | D17inj |
|---|---|---|---|
| D3inj | 0.05081 | - | - |
| D17inj | 0.00017 | 0.13482 | - |
| D31inj | 0.00022 | 0.42351 | 0.42351 |

**C. Comparing number of vacuoles in male brains of sham, D3Inj, D17Inj, and D31Inj conditions.**

Kruskal-Wallis rank sum test

data: total.no by injury
Kruskal-Wallis chi-squared = 15.336, df = 3, p-value = 0.001551

Pairwise comparisons using Wilcoxon rank sum test with continuity correction data: data.M$total.no and data.M$injury

| | sham | D3inj | D17inj |
|---|---|---|---|
| D3inj | 0.18399 | - | - |
| D17inj | 0.46556 | 0.79596 | - |
| D31inj | 0.00048 | 0.18399 | 0.18399 |

**D. Comparing total size of vacuoles in male brains of sham, D3Inj, D17Inj, and D31Inj conditions.**

Kruskal-Wallis rank sum test

data: total.area by injury
Kruskal-Wallis chi-squared = 15.163, df = 3, p-value = 0.001682

Pairwise comparisons using Wilcoxon rank sum test with continuity correction

data: data.M$total.area and data.M$injury

| | sham | D3inj | D17inj |
|---|---|---|---|
| D3inj | 1.0000 | - | - |
| D17inj | 1.0000 | 1.0000 | - |
| D31inj | 0.0029 | 0.0087 | 0.2101 |

### Males

| FlyBase ID | Gene name | Function | Log2FoldChange | Shared? |
|---|---|---|---|---|
| FBgn0039682 | Obp99c | | -0.774453287 | |
| FBgn0002565 | Lsp2 | energy storage | -0.932173432 | |
| FBgn0037146 | P5CS | energy metab | -0.76684635 | |
| FBgn0266347 | nAChRα4 | sleep promoting | 0.41036578 | |
| FBgn0033820 | CG4716 | no clue | -0.497357371 | |
| FBgn0025454 | Cyp6g1 | response to insecticide | -0.699296146 | |
| FBgn0028526 | CG15293 | head | -0.673721166 | |
| FBgn0000053 | Gart | IMP synthesis | -0.551393978 | w/ wt-mated females |
| FBgn0040653 | Dso1 | immune | -0.72422004 | |
| FBgn0001114 | Glt | axon outgrowth | -0.528692304 | |
| FBgn0001208 | Hn | serotonin and dopamine | -0.676318827 | |
| FBgn0267910 | lncRNA:CR34335 | | 0.359752339 | |
| FBgn0038147 | CCHa2 | stimulate food intake via horn | -0.449930577 | w/ wt-mated females |
| FBgn0034229 | CtsK2 | sexual reproduction | -0.779998723 | w/ wt-mated females |
| FBgn0034290 | CG5773 | visual | -1.100131223 | |
| FBgn0034885 | Eglp4 | renal system | -0.645754596 | |
| FBgn0038074 | Gnmt | metab | -1.578087349 | |
| FBgn0035090 | CG2736 | | -0.632428057 | |
| FBgn0015575 | α-Est7 | lipid metab | -0.568141504 | |
| FBti0062593 | transposable element | | 0.853653524 | |
| FBgn0034331 | BomBc2 | defense | -0.68335982 | |
| FBgn0263476 | snoRNA:CG32479-b | | 0.887811658 | |
| FBgn0259682 | Jabba | lipid droplet protein/ immune | -0.480897775 | |
| FBgn0025583 | BomS2 | imune | -0.584806198 | |
| FBgn0020385 | pug | energy metab | -0.591697635 | |
| FBgn0051869 | CG31869 | BMP binding | 0.29348842 | |
| FBgn0032726 | CG10621 | energy metab | -0.804015515 | w/ wt-mated females |
| FBgn0263461 | snoRNA:CG32479-a | | 0.802399758 | |
| FBgn0034162 | CG6426 | lysosome and sexual reproduc | -0.721540481 | |
| FBgn0000406 | Cyt-b5-r | maybe metab | -0.687927499 | |
| FBgn0020513 | Paics | IMP synthesis | -0.385932511 | w/ wt-mated females |
| FBgn0031561 | IM33 | immune | -0.623540942 | |
| FBgn0034200 | Gbp2 | response to stimulus | -0.574618097 | |
| FBgn0029823 | Shmt | metabolism and circadian rhy | -0.334951645 | |
| FBgn0041337 | Cyp309a2 | metab of insect hormones an | -0.96825435 | |
| FBgn0086691 | UK114 | protein chaperon similar to H | -0.761220024 | |
| FBti0019429 | transposable element | | 0.588459313 | |
| FBgn0032285 | CG17108 | head | -0.62564392 | w/ wt-mated females |
| FBgn0086687 | Desat1 | male mating behavior | -0.209450155 | w/ wt-mated females |
| FBgn0085195 | CG34166 | head | -0.593485014 | w/ wt-mated females |
| FBgn0261575 | tobi | carbo metab | -0.860328123 | |
| FBti0062383 | transposable element | | -1.899141011 | |
| FBgn0015039 | Cyp9b2 | metab of insect hormones an | -0.69275031 | |
| FBgn0087002 | apolpp | metab | -0.214002572 | w/ wt-mated females |
| FBgn0029828 | CG6067 | head | -0.69531812 | |
| FBgn0283427 | FASN1 | metab | -0.234710707 | w/ wt-mated females |
| FBgn0040736 | BomS3 | immune | -0.846028822 | |
| FBgn0029831 | CG5966 | metab | -0.903749038 | |
| FBgn0038465 | Irc | response to oxidative stress | -0.194903383 | |
| FBgn0058469 | lncRNA:CR40469 | | 0.441745013 | |
| FBgn0051769 | CG31769 | head | -0.661231957 | |
| FBgn0086670 | snoRNA:Ψ28S-2622 | | 1.015876829 | |
| FBgn0263470 | snoRNA:Tudor-SN-a | | 0.359047707 | |
| FBgn0039800 | Npc2g | sterol transport | -0.605902569 | |
| FBgn0033913 | CG8468 | metab | -0.690566207 | |
| FBgn0029990 | CG2233 | head | -0.653061653 | |
| FBgn0031449 | CG31689 | ATP binding | -0.276929092 | |
| FBgn0037191 | CG14448 | spermatozoon | 0.678817863 | |
| FBgn0010222 | Nmdmc | metab | -0.468618238 | |
| FBgn0259715 | CG42369 | | -0.465146308 | |
| FBgn0065073 | snoRNA:229 | | 1.288624169 | |
| FBgn0287606 | lncRNA:CR46475 | | 0.630041256 | |
| FBgn0040827 | CG13315 | | -0.401981202 | |
| FBgn0037387 | CG1213 | head | -0.414584191 | w/ wt-mated females |
| FBgn0029820 | CG16721 | signaling | -0.224444784 | |
| FBgn0039241 | CG11089 | wound healing | -0.417155457 | w/ wt-mated females |
| FBti0059668 | transposable element | | 0.749565983 | |
| FBgn0013763 | Idgf6 | chitin binding | -0.339309572 | w/ wt-mated females |
| FBti0020280 | transposable element | | 0.656605379 | |
| FBgn0031432 | Cyp309a1 | metab of insect hormones an | -0.855526769 | |
| FBgn0010383 | Cyp18a1 | | -0.794921059 | |
| FBgn0040398 | CG14629 | head | -0.420623202 | w/ wt-mated females |

### Females mated w/ wildtype males

| FlyBase ID | Gene name | Function | Log2FoldChange | Shared? |
|---|---|---|---|---|
| FBgn0038914 | fit | | -1.538791998 | |
| FBgn0032136 | Apoltp | lipid transport | -0.456213359 | |
| FBgn0030041 | CG12116 | | -0.93953107 | w/ SP0-mated females |
| FBgn0040398 | CG14629 | | -0.627198739 | w/ males |
| FBgn0283427 | FASN1 | energy | -0.449562538 | w/ males |
| FBgn0034394 | CG15096 | | -0.943667622 | |
| FBgn0034512 | Bbd | immune | -0.8244089 | |
| FBgn0032945 | CG8665 | | -1.461479676 | |
| FBgn0035091 | CG3829 | | -0.887598996 | |
| FBgn0003731 | Egfr | | -0.595581697 | |
| FBgn0032322 | CG16743 | | -0.477538657 | w/ SP0-mated females |
| FBgn0035770 | pst | LTM | -0.410712646 | |
| FBgn0086687 | Desat1 | immune | -0.399546646 | w/ males |
| FBgn0020415 | Idgf2 | apoptosis | -0.53096214 | |
| FBgn0029769 | frma | female receptivity | -1.124427796 | |
| FBgn0031461 | daw | hormone and meta | -0.579938076 | |
| FBgn0020513 | Paics | IMP synthesis | -0.604582893 | w/ males |
| FBgn0028371 | jbug | actin-binding and v | -0.715058385 | |
| FBgn0031538 | CG3246 | | -0.826485525 | |
| FBgn0051326 | CG31326 | | -1.115040157 | |
| FBgn0036501 | CG7272 | | -0.588751801 | |
| FBgn0013763 | Idgf6 | chitin-binding | -0.545141104 | w/ males |
| FBgn0011722 | Tig | hemolymph clot? | -0.619497397 | w/ virgins |
| FBgn0028543 | NimB2 | development | -0.526804832 | |
| RR46358_transposable_element | | | 0.517865925 | |
| FBgn0031560 | CG16713 | | -1.34235222 | |
| FBgn0043783 | CG32444 | | -0.960435429 | |
| FBgn0000053 | Gart | metabolism | -0.529627643 | w/ males |
| FBgn0026576 | Pisd | metabolism | -0.339393537 | |
| FBgn0034761 | CG4250 | | -0.430955497 | |
| FBgn0035978 | UGP | hyperoxia and met | -0.376705368 | |
| FBgn0264494 | CG17646 | metabolism | -0.331003508 | |
| FBgn0016122 | Acer | circadian rhythm a | -0.508768824 | |
| FBgn0032726 | CG10621 | maybe metab | -0.972836987 | w/ males |
| FBgn0261588 | pdm3 | smell perception | 0.23810159 | |
| FBgn0033366 | Ance-4 | protein metab | -0.956596672 | |
| FBgn0000163 | Baz | egg development | -0.581876885 | |
| FBgn0011705 | rost | development gene | -1.062344555 | |
| FBgn0263072 | CG43347 | neg reg of transcri | 0.198432807 | |
| FBgn0000592 | Est-6 | reproduction | -1.64582297 | |
| FBgn0039801 | Npc2h | imune | -1.407898481 | |
| FBgn0259145 | CG42260 | some ion channel | 0.199828821 | |
| FBgn0052687 | CG32687 | signal transduction | -0.678298246 | |
| FBgn0031305 | Iris | some virus gene? | -0.82524592 | |
| FBgn0039241 | CG11089 | involved in wound | -0.598911827 | w/ males |
| FBgn0267665 | lncRNA:CR46003 | | 0.335763938 | |
| FBgn0260474 | CG30002 | metabolism | -1.801990135 | |
| FBgn0263077 | M7BP | filopodium assemb | -0.516406784 | |
| FBgn0041182 | Tep2 | imune | -0.342373483 | |
| FBgn0038250 | CG3505 | protein metab | -1.024484163 | |
| FBgn0031220 | CG4822 | some membrane p | -0.483744486 | |
| FBgn0031805 | Nepl4 | proteolysis | -1.189140041 | |
| FBti0063239 | transposable element | | 0.605095347 | |
| FBti0062461 | transposable element | | 0.290893309 | |
| FBgn0037387 | CG1213 | membrane transpo | -0.996907971 | w/ males |
| FBgn0000473 | Cyp6a2 | reponse to oxygen | -0.491964115 | |
| FBgn0034229 | CtsK2 | reproduction | -0.983235651 | w/ males |
| FBgn0020617 | Rx | gene expression in | 0.414050792 | |
| FBgn0020416 | Idgf1 | chitin binding and | -0.624800361 | |
| FBgn0038147 | CCHa2 | stimulate food inta | -0.612545572 | w/ males |
| FBti0020410 | transposable element | | 0.281769862 | |
| FBgn0053196 | dpy | epidermal-cuticle a | -0.515861706 | |
| FBgn0010651 | MFS14 | response to hypoxi | -0.347089659 | |
| FBgn0051778 | CG31778 | | -0.93828276 | |
| FBgn0000477 | DNaseII | apoptosis | -0.45836991 | |
| FBgn0031630 | CG15629 | | -0.632988051 | |
| FBgn0034335 | GstE1 | response to heat | -0.524717997 | |
| FBgn0053120 | CG33120 | head specific | -0.803505993 | |
| FBgn0039161 | CG13606 | head specific | -1.179969857 | |
| FBgn0034717 | CG5819 | | -0.951727875 | |
| FBgn0005612 | Sox14 | tf in neurogenesis | -0.452415216 | |
| FBgn0013949 | Elal | response to nicotin | -0.492555788 | |
| FBgn0031913 | CG5958 | visual | -0.624020071 | |
| RR50446_transposable_element | | | 0.319234632 | |
| FBgn0032507 | CG9377 | protein metab | 1.267602408 | |
| FBgn0037819 | Phyhd1 | glial | -0.348431752 | |
| FBgn0038984 | AdipoR | insulin signaling | -0.249382708 | |
| RR48376_transposable_element | | | 0.385817452 | |
| FBgn0037801 | CG3999 | metab | -0.860189041 | |
| FBgn0004784 | inaC | visual | -0.654048852 | |
| FBgn0001149 | GstD1 | not sure | -0.240434463 | |
| FBgn0023441 | fus | splicing factor | -0.782661332 | |

### Virgin females

| FlyBase ID | Gene name | Function | Log2FoldChange | Shared? |
|---|---|---|---|---|
| FBgn0036232 | CG14125 | chitin binding | -28.77174582 | |
| FBgn0022355 | Tsf1 | immune | 0.36142726 | |
| FBgn0032897 | bero | neg. reg. of pain | 0.659910491 | |
| FBgn0030159 | CG9689 | head | 0.651705339 | |
| FBgn0036422 | CG3868 | | -3.418196822 | |
| FBgn0085353 | CG34324 | chitin | -7.915609665 | w/ SP0-mated females |
| FBgn0085249 | CG34220 | chitin | -6.32508532 | w/ SP0-mated females |
| FBgn0267348 | LanB2 | development | 0.375229227 | |
| FBgn0043575 | PGRP-SC2 | immune | -1.904962432 | |
| FBgn0032899 | CG9338 | sleep | 0.604613976 | |
| FBgn0051901 | Mur29B | extracellular ma | -2.922528027 | |
| FBgn0002526 | LanA | development | 0.322748869 | |
| FBgn0036203 | Muc68D | extracellular ma | -9.171413766 | w/ SP0-mated females |
| FBgn0002578 | Kaz-m1 | not sure | -3.342784876 | |
| FBgn0014019 | Rh5 | visual | 0.83522137 | |
| FBgn0011722 | Tig | hemolyph clot | 0.674428238 | w/ wt-mated females |

### Females mated w/ SP0

| FlyBase ID | Gene name | Function | Log2FoldChange | Shared? |
|---|---|---|---|---|
| FBgn0004240 | DptA | immune | -7.102691081 | |
| FBgn0014859 | Hr38 | response to hor | -0.309375242 | |
| FBgn0085353 | CG34324 | chitin binding | 2.946951041 | w/ virgins |
| FBgn0085249 | CG34220 | chitin binding | 1.410945615 | w/ virgins |
| FBgn0034638 | CG10433 | receptivity | -0.272699497 | |
| FBgn0263776 | CG43693 | | -0.26749276 | |
| FBgn0036203 | Muc68D | chitim binding | 3.758412571 | w/ virgins |
| FBgn0041194 | Prat2 | IMP synthesis | -0.665311222 | |
| FBgn0261714 | Cpn | visual | -1.070638941 | |
| FBgn0030258 | CG1552 | | -0.224968211 | |
| FBgn0030041 | CG12116 | | -0.502028091 | w/ wt-mated females |
| FBgn0032322 | CG16743 | | -0.273258373 | w/ wt-mated females |
| FBgn0035880 | Culd | visual | -0.637988482 | |

| | | | | |
|---|---|---|---|---|
| FBgn0032285 | CG17108 | head | -1.145016404 | w/ males |
| FBgn0036975 | CG5618 | metab | -0.417414818 | |
| FBgn0014031 | Agxt | enzyme | -0.847162291 | |
| FBgn0035280 | Cpr62Bb | cuticle and chitin | -1.430096524 | |
| FBgn0051103 | CG31103 | transmembrane tra | -1.413030803 | |
| FBgn0264699 | CG43968 | | 1.469582938 | |
| FBgn0051538 | CG31538 | head | 0.281457134 | |
| FBgn0262139 | trh | trachea | -0.379669256 | |
| FBti0060955 | transposable element | | 1.085000568 | |
| FBgn0085195 | CG34166 | head | -0.94390951 | w/ males |
| FBgn0004047 | Yp3 | egg development | -0.866771324 | |
| FBgn0030266 | CG11122 | nervous system | 0.186775665 | |
| FBgn0035321 | CG1275 | nervous sytem | -0.210184915 | |
| FBgn0052702 | Cubn | required in nephro | 0.244266741 | |
| FBgn0039311 | CG10513 | Ecdysteroid kinase- | -0.397686723 | |
| FBgn0033170 | sPLA2 | metab | -0.778661531 | |
| FBgn0000003 | 7SLRNA:CR32 | targets ER | 0.318053478 | |
| FBgn0031107 | HERC2 | protein ubiquitinat | 0.214778025 | |
| FBti0062672 | transposable element | | 1.805123117 | |
| FBgn0030925 | Hayan | imune | -1.380572537 | |
| FBgn0026061 | Mipp1 | trachea developme | -0.558071845 | |
| FBgn0034432 | Acadvl | metab | -0.31762541 | |
| FBgn0087002 | apolpp | metab | -0.600273003 | w/ males |
| FBgn0005391 | Yp2 | egg dev | -1.006638368 | |
| FBgn0026403 | Ndg | cell adhesion glyco | -0.597137781 | |
| FBgn0033782 | sug | carbohydrate meta | -0.857540413 | |
| FBgn0004623 | Gβ76C | visual | -0.629920979 | |
| FBgn0036433 | CG9628 | septate junction as | -0.526279043 | |

### Males

| FlyBase ID | Gene name | Function | Log2FoldChange | Shared? |
|---|---|---|---|---|
| FBgn0000120 | Arr1 | Visual | -0.660051779 | |
| FBgn0004784 | inaC | Visual | -1.153972103 | |
| FBgn0041581 | AttB | immune | -2.940765803 | |
| FBgn0003250 | Rh4 | visual | -0.650760983 | |
| FBgn0026314 | Ugt35B1 | | -0.263185618 | |
| FBgn0001233 | HSP83 | immune | -0.41774832 | |
| FBgn0033728 | Cpr49Ae | cuticle | -2.916260656 | |
| FBgn0012042 | AttA | immune | -1.244612229 | |
| FBgn0002938 | ninaC | visual | -0.483038218 | |
| FBgn0034328 | BomBc1 | immune | -1.722311387 | |

### Females mated w/ wildtype males

| FlyBase ID | Gene name | Function | Log2FoldChange | Shared? |
|---|---|---|---|---|
| FBgn0014865 | Mtk | immune | -1.848555919 | |
| FBgn0030304 | Cyp4g15 | | -0.231873949 | |
| FBgn0034407 | DptB | immune | -6.216305169 | |
| FBgn0036262 | Miox | | 0.360308721 | |
| FBgn0038877 | CG3308 | | -0.302970343 | |
| FBgn0268063 | ltl | development | -0.288749888 | |
| FBgn0033367 | PPO2 | immune | 0.445630261 | |
| FBgn0001233 | HSP83 | immune | -0.243474805 | w/ SP0-mated females |

### Virgin females

| FlyBase ID | Gene name | Function | Log2FoldChange | Shared? |
|---|---|---|---|---|
| FBgn0029966 | Ir7c | Taste receptor | -4.790054545 | |
| FBti0020280 | | | 0.420139294 | |

### Females mated w/ SP0

| FlyBase ID | Gene name | Function | Log2FoldChange | Shared? |
|---|---|---|---|---|
| FBgn0026418 | Hsp110 | Response to heat shock | -0.319378612 | |
| FBgn0036145 | CG7607 | | 0.368973498 | |
| FBgn0037788 | CAH7 | metab | 0.289189632 | |
| FBgn0010225 | Gel | actin binding | 0.219140227 | |
| FBgn0012344 | Dh44 | hormones and response | -0.223383066 | |
| FBgn0015577 | α-Est9 | | 0.28242756 | |
| FBgn0034398 | CG15098 | | 0.203139536 | |
| FBgn0036663 | CG9674 | NADH activity | -0.168419631 | |
| FBgn0259108 | futsch | actin binding and axon re | -0.180246072 | |
| FBgn0262593 | Shab | neuronal excitability | -0.142266267 | |
| FBgn0039844 | CG1607 | | -0.248854791 | |
| FBgn0001233 | HSP83 | Immune | -0.528520138 | w/ wt-mated females |
| FBgn0004575 | Syn | synaptic vesicles | -0.145657451 | |
| FBgn0014906 | Hydr2 | lipid metab | 0.189310644 | |
| FBgn0031313 | CG5080 | | 0.201602684 | |
| FBgn0263106 | DnaJ-1 | HSP co-factor regulating | -0.564874198 | |
| FBgn0020618 | Rack1 | devleopment and reprod | 0.157630723 | |
| FBgn0086677 | jeb | dev | -0.221336404 | |
| FBgn0051221 | CG31221 | | -0.156178598 | |
